# Donor-resolved meta-programs identify a T2D-associated β-cell program linked to insulin secretory capacity in human islets

**DOI:** 10.64898/2026.11.23.768764

**Authors:** Yuelong Chen, Tengxiao Lei, Shiya Huang, Haokun Han, Yanci Wang, Jiawei Li, Lixin Cheng, Weidong Tang, Jijun Tang, Nana Jin

**Affiliations:** College of Electronics & Information Engineering, Jinggangshan University, Ji’an, China; School of Life Sciences, The Chinese University of Hong Kong, Hong Kong SAR, China; School of Computer Science and Artificial Intelligence, Shenzhen University of Advanced Technology, Shenzhen, Guangdong, China; Guangdong Provincial Clinical Research Center for Geriatrics, Shenzhen People’s Hospital (The Second Clinical Medical College, Jinan University; The First Affiliated Hospital, Southern University of Science and Technology), Shenzhen, Guangdong, China

**Keywords:** type 2 diabetes, pancreatic islets, β cell, single-cell RNA sequencing, meta-program, non-negative matrix factorization, insulin secretion, islet perifusion, transcriptional signature

## Abstract

Reproducible transcriptional programs that link type 2 diabetes (T2D) to human β-cell secretory dysfunction remain poorly defined. We applied donor-resolved non-negative matrix factorization to single-cell transcriptomes from 48 donors without diabetes, with prediabetes, or with T2D, identifying 43 recurrent meta-programs across seven islet cell types. Gene sets and scoring procedures were fixed before association testing and validation. A 50-gene β-cell program, β-MP5, showed the largest standardized reduction in T2D. This reduction was reproduced in the Human Pancreas Analysis Program cohort and pooled Smart-seq2 datasets, with directionally consistent effects across four bulk cohorts. In donor-matched perifusion measurements from two laboratories with overlapping donors, higher β-MP5 scores were associated with greater insulin secretion under glucose stimulation, cAMP amplification, and membrane depolarization. These associations persisted after adjustment for diagnosis, islet viability, β-cell fraction, and cold ischemia time. Spatial transcriptomics further showed higher β-MP5 scores in islet-rich regions of intact pancreas. Together, these findings identify a recurrent β-cell transcriptional program that is reduced in T2D and tracks ex vivo insulin secretion, providing a fixed molecular readout for investigating human β-cell functional states.

## Introduction

Diabetes affects more than half a billion adults worldwide, most of whom have type 2 diabetes (T2D), and its prevalence continues to rise ^1^. Although insulin resistance is common, overt hyperglycaemia develops when pancreatic β cells can no longer secrete sufficient insulin to meet metabolic demand ^2,3^. Both a reduction in β-cell mass and a decline in the secretory capacity of the remaining β cells contribute to this failure ^4–6^, and islets isolated from donors with T2D show impaired glucose-stimulated insulin secretion ex vivo ^7^. Several mechanisms have been proposed to underlie β-cell dysfunction, including glucolipotoxicity, oxidative stress and hypoxia, loss of key β-cell transcription factors and β-cell dedifferentiation ^8–13^. Importantly, β-cell function can recover in some individuals after substantial weight loss in the early years after diagnosis ^14^, suggesting that at least part of the defect reflects a potentially reversible change in β-cell state. Molecular readouts that capture these changes in β-cell state and relate them to functional capacity are therefore of considerable translational interest.

Transcriptomic profiling of human islets has been used extensively to identify molecular features associated with β-cell dysfunction. Bulk studies of islets from organ donors and of laser-capture microdissected islets from living donors undergoing pancreatectomy have identified T2D-associated genes and co-expression modules related to insulin secretion and glycaemia ^15–18^. However, bulk expression profiles combine signals from endocrine and exocrine cells and are influenced by differences in donor characteristics, islet procurement and culture conditions ^19,20^. Single-cell RNA sequencing (scRNA-seq) has resolved the cellular composition of human islets ^21–27^, yet the T2D-associated gene expression changes reported for β cells have shown limited overlap across studies ^28–31^. Differences in donor numbers, technical platforms and analytical strategies, including approaches that do not adequately account for donor-level replication, have complicated comparisons across studies ^32,33^. Larger single-cell cohorts and resources that combine islet transcriptomes with physiological phenotyping, such as the Human Pancreas Analysis Program (HPAP), now provide opportunities to investigate these changes at the donor level ^34–38^.

β cells are also heterogeneous within an individual. Studies of human and mouse islets have described β-cell subpopulations and states that differ in maturity, proliferative capacity, stress responses and secretory behaviour ^25,36,39–41^. Because such variation is often continuous and individual cells can participate in multiple biological processes simultaneously, it may be better represented by gene expression programs than by discrete cell clusters. Non-negative matrix factorization (NMF) decomposes expression data into additive programs of co-varying genes ^42,43^. In cancer, applying NMF separately to individual tumours and subsequently clustering recurrent programs across tumours has identified consensus meta-programs (MPs) that capture both generic and lineage-specific cellular processes ^44–46^. A similar framework has recently been applied to cell types in the Alzheimer’s disease brain ^47^. By decomposing each sample separately, this approach retains the individual as the unit of discovery and identifies programs that recur across individuals, providing a framework for characterizing shared transcriptional states in human tissues with substantial donor-to-donor variation.

However, to our knowledge, this framework has not been systematically applied to human islets in T2D. It therefore remains unclear whether recurrent transcriptional programs across islet cell types are altered in T2D, whether these alterations are reproducible across independent cohorts and measurement platforms, and whether they are related to measured β-cell function. Establishing such functional relationships is particularly important because disease-associated gene expression differences do not necessarily translate into altered cellular function. Islet transcriptomes have so far been linked to secretory measurements mainly at the level of single cells or β-cell subtypes ^36,48^, and relating donor-level transcriptional programs to whole-islet insulin secretion could provide a more direct view of the molecular states associated with human β-cell dysfunction. We hypothesized that donor-resolved MPs would reveal T2D-associated programs that are reproducible across cohorts and related to measured β-cell function, rather than simply reflecting diagnostic status.

In this study, we applied donor-resolved NMF to single-cell transcriptomic data from 48 donors without diabetes, with prediabetes or with T2D and identified 43 recurrent MPs across seven major islet cell types. We fixed the MP gene sets and scoring procedure before any disease-association testing or validation analyses and then evaluated their associations with T2D and HbA1c. Among all programs, the β-cell program β-MP5 showed the most marked reduction in T2D. We evaluated its reproducibility in an independent single-cell cohort, in pooled Smart-seq2 datasets and across five bulk islet transcriptomic cohorts. We further related β-MP5 scores to donor-matched islet perifusion measurements from two laboratories and examined its spatial distribution in human pancreatic tissue. Together, these analyses identify β-MP5 as a reproducible, T2D-associated β-cell transcriptional program that tracks ex vivo insulin secretory function in human islets.

## Methods

### Study design and data sources

This study used publicly available or access-controlled data from previously published studies; no new human samples were collected. The analysis comprised a discovery stage followed by independent validation and functional analyses. The discovery cohort was the single-cell RNA-sequencing (scRNA-seq) dataset GSE221156, comprising islets from 48 organ donors without diabetes (ND, n = 17), with prediabetes (PD, n = 14) or with type 2 diabetes (T2D, n = 17) ^37^.

Validation data comprised (i) islet scRNA-seq data from the Human Pancreas Analysis Program (HPAP), accessed through PanKbase/PANC-DB ^34,35^; (ii) three Smart-seq2 islet scRNA-seq datasets (GSE153855, E-MTAB-5061 and GSE214517) ^23,38^; (iii) five bulk islet transcriptomic datasets (GSE76894, GSE164416, GSE20966, GSE25724 and GSE50244) ^15–18^; (iv) islet perifusion and donor phenotype data for HPAP donors obtained through PANC-DB; and (v) 10x Genomics Visium spatial transcriptomic data from human pancreas sections (GSE264331) ^49^. Validation datasets were not used for meta-program discovery.

The meta-program (MP) gene sets, scoring procedure, gene backgrounds and planned disease contrasts were fixed and saved with checksums on 26 October 2026, before disease-association testing and validation analyses. The frozen gene sets and scoring objects were maintained unchanged throughout subsequent analyses.

For each dataset, donors were registered using their original identifiers, diagnosis, age, sex, body mass index (BMI) and HbA1c, where available. Missing clinical values were not imputed. Potential donor overlap between datasets was assessed using shared identifiers, published sample descriptions and, where identifiers were unavailable, comparisons of sex, age and BMI. One GSE214517 donor identified as overlapping with the discovery cohort and three additional GSE214517 donors whose identity with E-MTAB-5061 donors could not be excluded were removed from the Smart-seq2 validation. HPAP donors were independent of the discovery cohort ^37^. For bulk datasets originating from the same procurement centre, sample identifiers were compared and potential overlap was documented (Table S7).

**Ethics statement.** This study used publicly available or access-controlled, de-identified data from previously published studies. No new human samples were collected. The original studies obtained informed consent and ethics approval from their respective institutions. The present analyses were conducted in accordance with the access conditions of the source datasets.

### Quality control and cell-type annotation

Filtered UMI count matrices for the 54 GSE221156 libraries were obtained from the Gene Expression Omnibus (GEO). Cells were assigned to donors using the cell-to-donor mapping table provided by the original study ^37^. Cells not retained by the original authors were not assigned to donors.

Quality control (QC) was performed separately for each capture library. Doublets were identified using scDblFinder ^50^, with an expected doublet rate of 0.8% per 1,000 cells, and only singlets were retained. Cells were subsequently required to have at least 500 UMIs, at least 250 detected genes and a mitochondrial transcript fraction of no more than 0.40. Additional outliers were removed if their log-transformed UMI counts or detected gene counts were more than three median absolute deviations (MADs) below the library median, or if their mitochondrial transcript fraction was more than three MADs above the library median.

Cell-type annotation was performed using Seurat ^51^. Counts were log-normalized with a scale factor of 10,000, and 2,000 variable genes were selected per library. Principal component analysis (PCA; 30 components) was followed by Harmony integration across libraries ^52^, graph-based Louvain clustering (resolution = 0.8) and UMAP embedding ^53^. The annotation embedding also included cells from two additional islet datasets, which were used exclusively to support marker-panel-based clustering and were excluded from all downstream analyses.

Each cell was scored using UCell ^54^ for predefined marker panels representing β cells (INS, IAPP, MAFA, NPTX2, HADH and DLK1), α cells (GCG, ARX, IRX2, TTR and LOXL4), δ cells (SST, HHEX and LEPR), γ cells (PPY, ETV1 and CARTPT), ductal cells (KRT19, CFTR, SOX9 and KRT7), acinar cells (PRSS1, CPA1, CELA3A and CTRB2), stellate cells (COL1A1, PDGFRB, RGS5 and SPARC), endothelial cells (PECAM1, PLVAP and VWF), immune cells (PTPRC, CD68, LAPTM5 and TYROBP) and mast cells (TPSAB1, CPA3 and KIT).

A marker panel was considered eligible for a cluster when at least two of its genes were detected in at least 20% of the cluster’s cells. Each cell voted for its highest-scoring eligible panel. A cluster was assigned a cell-type label when more than 50% of its cells voted for the same panel and no other panel received 20% or more of the votes. Clusters with competing hormone-panel assignments were classified as multihormonal, while other unresolved clusters were classified as uncertain. Cluster-level assignments were additionally required to agree with an unintegrated donor-level pseudobulk vote. Hormone transcripts alone were not used to determine cell identity, and disease labels were not considered during annotation. Agreement between the final labels and the original authors’ annotations was assessed descriptively. Integrated embeddings were used only for annotation and visualization; all subsequent analyses used unintegrated expression counts.

### Donor-resolved non-negative matrix factorization

Non-negative matrix factorization (NMF) was performed separately for each donor and cell type, following the meta-program discovery framework described by Gavish et al. ^46^. Each donor × cell-type unit containing at least 10 cells of one of the seven principal cell types was analysed. This yielded 322 units: 48 donors for α, β, δ, ductal, acinar and stellate cells, and 34 donors for γ cells.

Within each unit, raw counts were converted to counts per million (CPM), and the 7,000 genes with the highest mean expression were retained. Expression values were transformed as log2(CPM/10 + 1), centred across cells for each gene, and negative values were set to zero. No cell subsampling was performed.

NMF was performed using the snmf/r algorithm implemented in the R package NMF ^55^. Ranks from 4 to 9 were evaluated, with 10 runs per rank using a fixed random seed. Each factor was represented by its 50 highest-loading genes. Across the 322 donor × cell-type units, this procedure yielded 39 candidate programs per unit and 12,558 candidate programs in total.

Robust programs were selected using the criteria and implementation described by Gavish et al. ^46^. A program was retained if it shared at least 35 of its 50 genes with a program obtained at another NMF rank within the same donor and cell type, shared at least 10 genes with a program from another donor of the same cell type, and was not redundant with a previously selected program from the same donor × cell-type unit. Redundancy was defined as an overlap of more than 10 genes during greedy selection. This procedure yielded 1,799 robust programs.

### Meta-program construction

Within each cell type, robust programs were clustered into MPs using the greedy clustering algorithm described by Gavish et al. ^46^, based on the number of shared genes between programs.

Clustering was initiated with the program having the largest number of programs sharing at least 10 genes with it; at least six such programs were required to initiate a cluster. Additional programs were incorporated while they shared at least 10 genes with the current MP gene set. Each MP was represented by the 50 genes occurring most frequently among its member programs. Ties at the 50th position were resolved using NMF loading. All 43 resulting MPs were retained without manual selection, and MPs were numbered separately within each cell type.

Independent donor support for each MP was calculated as the number of distinct donors contributing at least one member program. Each donor was counted only once per MP, irrespective of the number of programs contributed.

### Functional annotation of meta-programs

Over-representation analyses were performed using MSigDB Hallmark, KEGG (legacy) and Gene Ontology biological process (GO:BP) gene sets (MSigDB release 2026.1.Hs) ^56–58^. Enrichment was tested separately for each MP using the hypergeometric test implemented in clusterProfiler ^59^.

For each cell type, the gene universe was defined as the union of genes entering NMF across its donor × cell-type units. Gene sets containing 10–500 genes within the tested universe were retained. P values were adjusted using the Benjamini–Hochberg (BH) procedure ^60^, separately within each MP and gene-set collection. Enrichment was considered significant at FDR < 0.05.

### Meta-program scoring

MPs were scored using a fixed rank-based procedure applied to donor-level pseudobulk expression profiles. For each donor and cell type, raw UMI counts from all QC-passed cells of the corresponding cell type were summed to generate a pseudobulk profile.

Within each pseudobulk sample, genes in a predefined background set (B) were ranked by expression, with tied values assigned their average rank.

The MP score was calculated as the mean of \(2q_g-1\) across all member genes of the MP present in the background. The background consisted of genes in the frozen gene universe that were measured in the respective dataset, defined from the dataset’s feature list without reference to disease outcomes.

A score was calculated only when at least 80% of the 50 MP genes and at least 10 genes were represented in the background. In the discovery cohort, donor × cell-type pseudobulk profiles used for disease-association analyses required at least 30 cells. For gene-expression heatmaps, pseudobulk counts were normalized using the trimmed mean of M-values (TMM) method and expressed as log-counts per million (log-CPM) using edgeR ^61^. Each gene was standardized across donors before group-level visualization.

### Disease associations in the discovery cohort

MP scores were compared between T2D and ND donors and between PD and ND donors using ordinary least-squares linear regression adjusted for age, sex and BMI. Each diagnostic contrast was fitted separately using only the two groups being compared. Associations with HbA1c (%) were estimated across all 48 donors using the same covariates.

Heteroskedasticity-consistent HC3 standard errors were used for inference ^62–64^. Diagnostic effects and HbA1c associations were expressed in units of the standard deviation (SD) of MP scores among ND donors. A model was considered estimable when each diagnostic group contained at least five donors and the residual degrees of freedom were at least five.

The false discovery rate (FDR) was controlled using the BH procedure within prespecified testing families. For diagnostic comparisons, T2D-versus-ND and PD-versus-ND tests were corrected together within each cell type; for β-cell programs, this family comprised 16 tests. For HbA1c associations, all 43 MPs were included in a single testing family. For β-MP5 and β-MP3, leave-one-donor-out analyses were performed by refitting the T2D-versus-ND model after removing each ND or T2D donor in turn. Standardization continued to use the ND SD from the corresponding full analysis.

### Independent single-cell validation in HPAP

HPAP islet scRNA-seq data were obtained from PanKbase as STARsolo-generated raw UMI count matrices using GENCODE version 39. Cell calling was performed using EmptyDrops, with a lower UMI bound of 100, 100,000 iterations and FDR < 0.005. The QC thresholds and cell-type annotation procedure used in the discovery cohort were subsequently applied to HPAP. Harmony-integrated clustering was used for cell annotation only.

Of the 83 HPAP donors with islet scRNA-seq data, donors with type 1 diabetes (n = 12), autoantibody-positive status without diabetes (n = 12) and one donor with only treated samples available were excluded. The resulting validation cohort comprised 58 donors: 36 ND and 22 T2D.

Donor-level β-cell pseudobulk profiles were generated and scored using the frozen MP signatures and the HPAP-specific gene background. The primary validation analysis included donors with at least 30 profiled β cells, yielding 55 donors (34 ND and 21 T2D). Scores were compared between T2D and ND donors using HC3 regression adjusted for age, sex, BMI and 10x chemistry (v2 or v3). Chemistry information was obtained from PanKbase records or, for four donors, inferred by matching cell barcodes to the 10x v2 and v3 barcode whitelists. Effects were expressed in units of the ND donor SD, and FDR was controlled across the eight β-cell MPs.

For gene-level concordance, β-cell pseudobulk counts from the discovery and HPAP cohorts were analysed using the edgeR quasi-likelihood framework ^65^. Genes passing filterByExpr were retained together with all β-MP5 member genes. Libraries were TMM-normalized, and T2D-versus-ND differences were estimated using glmQLFit followed by the quasi-likelihood F-test, with robust dispersion and prior estimation. Models were adjusted for age, sex and BMI and, in HPAP, 10x chemistry. Agreement between cohorts was assessed using the Spearman correlation of T2D-versus-ND log2 fold changes across the 50 β-MP5 genes and, for reference, across all genes tested in both cohorts.

The β-cell-only HPAP UMAP shown in Figure 4A was calculated using 2,000 variable genes and 20 principal components without batch integration.

### Pooled Smart-seq2 validation

Three additional human islet scRNA-seq datasets generated using Smart-seq2 were analysed: GSE153855, E-MTAB-5061 and GSE214517. Per-sample read counts were obtained for GSE153855 and E-MTAB-5061. For GSE214517, read counts were regenerated from FASTQ files using STARsolo in Smart-seq mode because only RPKM values were available in the deposited dataset.

The three datasets were processed using the same cell-type annotation procedure. Cells were required to have at least 50,000 reads and at least 1,000 detected genes. For GSE214517, an additional mitochondrial transcript fraction threshold of 0.30 or less was applied.

Because the three datasets originated from the same research group and had relatively small individual sample sizes, they were analysed jointly. The pooled dataset comprised 27 donors and 7,254 QC-passed cells. MP scores were calculated from donor-level β-cell pseudobulk profiles using the intersection of the three dataset gene backgrounds (19,494 genes, including 47 of the 50 β-MP5 genes). Donors with at least 30 β cells were included, yielding 19 donors (11 ND and eight T2D).

Scores were compared between T2D and ND donors using HC3 regression adjusted for dataset. Effects were expressed in units of the SD of dataset-centred ND scores, and FDR was controlled across the eight β-cell MPs. Robustness was assessed using leave-one-dataset-out analyses and, among the 11 donors with complete clinical data, additional adjustment for age, sex and BMI.

### Bulk islet transcriptomic cohorts

Five bulk islet transcriptomic datasets were analysed: GSE76894, GSE164416, GSE20966, GSE25724 and GSE50244.

Affymetrix microarray datasets (GSE76894, GPL570; GSE20966, GPL1352; GSE25724, GPL96) were processed from raw CEL files using robust multi-array average (RMA) normalization ^66^, with platform-specific CDF and annotation packages. Probe sets mapping to multiple genes were removed. Where multiple probe sets mapped to the same gene, the probe set with the highest mean RMA-normalized expression across samples was retained.

For GSE164416, HTSeq gene-level counts were TMM-normalized and converted to log-CPM. For GSE50244, the deposited normalized expression matrix was used. Gene symbols were mapped one-to-one to Ensembl identifiers using current HGNC symbols, followed by previous symbols and aliases.

The frozen β-MP5 signature was scored separately within each cohort using that cohort’s own measured gene background. GSE25724 measured 38 of the 50 β-MP5 genes (76%), below the 80% coverage criterion. It was therefore included only as a labelled sensitivity analysis, with the coverage requirement relaxed for this dataset.

T2D-versus-ND differences in MP scores were estimated using HC3 regression adjusted for age, sex and BMI and, where applicable, processing batch. Processing batch was obtained from GEO annotations for GSE76894 and from library preparation date for GSE164416. Batches containing only one donor were excluded. For GSE164416, sex information was obtained from matching patient records in GSE76895. The primary diagnostic and HbA1c analyses in GSE164416 included ND and T2D donors. Donors with impaired fasting glucose or impaired glucose tolerance were additionally included in a sensitivity analysis, while donors with type 3c diabetes were excluded.

For GSE50244, which lacked diagnostic group information, associations between β-MP5 scores and HbA1c were estimated using HC3 regression adjusted for age, sex and BMI. Effects were expressed in units of the ND donor SD, or the SD across all donors for GSE50244, which had no diagnostic groups.

Gene-level T2D-versus-ND differences were estimated separately within each cohort using the covariates included in the corresponding MP score model. Limma with empirical Bayes moderation (trend = TRUE) was used for microarray datasets ^67^, and the edgeR quasi-likelihood framework was used for GSE164416 ^65^. Genome-wide FDR was calculated across all genes tested within each cohort.

### Bulk cell-type deconvolution

Cell-type composition was estimated using BayesPrism ^68^, MuSiC ^28^ and CIBERSORTx ^69^. A reference expression profile was constructed from the discovery cohort using QC-passed cells belonging to the seven principal islet cell types, with up to 100 cells per donor and cell type for the BayesPrism and MuSiC reference and up to 300 cells per cell type for the CIBERSORTx signature matrix.

BayesPrism was run with default settings after removing ribosomal, mitochondrial, sex-chromosome and MALAT1 genes and restricting the reference to protein-coding genes. MuSiC was run using weighted proportion estimates. CIBERSORTx was run in relative mode with S-mode batch correction and 100 permutations.

Because estimates differed between methods and were not interpretable across all cohorts, cell-composition estimates were reported descriptively and were not included as covariates in the primary bulk analyses. This decision reflected the limited consistency and applicability of deconvolution estimates across platforms and tissue preparations. The potential contribution of β-cell fraction to bulk expression profiles is considered in the interpretation of the results.

### Islet perifusion and functional measures

Islet perifusion data and associated donor phenotype information for HPAP donors were obtained from PANC-DB. Measurements were available from two laboratories: the University of Pennsylvania (UPenn) and Vanderbilt University.

At UPenn, insulin release was measured in 1-min fractions and expressed as ng per 100 islets per minute. Islets were perifused sequentially with 3 mM glucose, 16.7 mM glucose, 16.7 mM glucose plus 0.1 mM IBMX and 30 mM KCl. Most preparations were studied in the presence of a background mixture of amino acids at 4 mM.

At Vanderbilt, insulin release was measured in 3-min fractions and expressed as ng per 100 islet equivalents per minute. Perifusion was performed on a background of 5.6 mM glucose, with sequential stimulation using 16.7 mM glucose, 16.7 mM glucose plus 100 µM IBMX and 20 mM KCl ^19^.

For each stimulation phase, the insulin secretion area under the curve (AUC) was calculated as the mean insulin release across the phase multiplied by the phase duration, without baseline subtraction. An AUC was set to missing if fewer than 80% of the fractions within the corresponding phase were measured.

Additional functional outcomes included islet insulin content, the UPenn glucose stimulation index (mean insulin release at 16.7 mM divided by mean release at 3 mM glucose), the PANC-DB glucose-stimulated insulin response (GSIR) index and donor serum C-peptide. All functional outcomes were log-transformed before regression analysis.

The predictor was the β-MP5 score calculated from each donor’s β-cell pseudobulk profile using the frozen scoring procedure. Scores were standardized using the mean and SD of HPAP ND donors in the corresponding analysis set, consistent with the HPAP validation analysis.

The primary functional analyses included donors with at least 30 profiled β cells (UPenn, n = 52; Vanderbilt, n = 42). Thirty-nine donors had measurements available from both laboratories. The two laboratory datasets were analysed separately because of differences in perifusion protocols, measurement units and donor overlap; raw AUC values were not pooled.

Associations between β-MP5 scores and functional outcomes were estimated using HC3 linear regression with three nested models:

– Model 1: adjusted for age, sex, BMI and 10x chemistry.
– Model 2: Model 1 with additional adjustment for diagnosis.
– Model 3: Model 2 with additional adjustment for islet viability, β-cell fraction and cold ischaemia time.

β-cell fraction was calculated as the number of β cells divided by the total number of α, β, δ and γ cells in the scRNA-seq data. Complete cases were used for each model. A model was considered estimable when at least 10 donors were included, at least five residual degrees of freedom remained and no model terms were aliased.

The primary FDR family comprised the β-MP5 associations with the three perifusion AUC outcomes and insulin content in Model 1, separately for each laboratory. The same outcomes were evaluated for the other seven β-cell MPs as a separate testing family. Secondary functional outcomes and the additional adjustment models were evaluated in separate testing families. FDR was controlled using the BH procedure, and FDR < 0.05 was considered statistically significant; for the other seven β-cell MPs, we additionally report whether any association reached FDR < 0.1. Influence was assessed using leave-one-donor-out estimates and Cook’s distance.

### Spatial transcriptomics

10x Genomics Visium spatial transcriptomic data from 16 whole-pancreas sections obtained from six donors (three with normal glucose tolerance [NGT] and three with T2D) were downloaded from GEO (GSE264331) ^49^. Raw feature–barcode matrices were restricted to in-tissue spots, and donor identities were assigned using the published sample tables and checked against sex-chromosome gene expression. Spots with fewer than 500 UMIs were excluded.

Because hormone transcripts were subject to substantial spillover into adjacent tissue in this dataset ^49^, islet-rich spots were identified using seven non-hormone endocrine marker genes: CHGA, CHGB, SCG2, SCG3, SCG5, PCSK1N and PCSK2. None of these genes was a member of β-MP5.

For each spot, the expected number of marker UMIs was calculated as the total UMI count multiplied by the section-wide marker fraction. Spots containing at least three marker UMIs and exceeding the 99.9th percentile of the corresponding Poisson distribution were classified as islet-rich. Spots with marker counts at or below the expected count were classified as endocrine-low, while intermediate spots were excluded.

The frozen β-MP5 scoring procedure and the Visium-specific gene background were applied to individual spots for spatial visualization and to donor-level pseudobulk profiles for statistical testing. For donor-level analyses, counts from islet-rich and endocrine-low spots were summed separately across all available sections for each donor. Within-donor differences between the two compartments were assessed using an exact two-sided sign-flip test across all 2⁶ possible sign combinations. Donors, rather than individual spots or sections, were the units of inference.

For visualization, spot scores were averaged with the scores of the six adjacent spots. Colour scales were clipped at the second and 98th percentiles within each section. Because sequencing depth differed substantially between sections, colour intensities were interpreted within sections rather than used for direct quantitative comparisons between donors.

### Statistical analysis and software

Donors were the unit of inference throughout the study. Cells, spots, sections and technical replicates were aggregated at the donor level before inferential testing, except for analyses explicitly describing spatial distributions. All statistical tests were two-sided.

Unless otherwise specified, linear regression models used HC3 heteroskedasticity-consistent standard errors. Effects are reported with 95% confidence intervals. Multiple testing was controlled using the BH procedure within the testing families defined for each analysis. Missing clinical and functional measurements were not imputed; analyses used the donors with complete data for the variables included in the corresponding model.

Analyses were performed in a Docker environment based on Bioconductor 3.23 and R 4.6.1. The principal R packages included Seurat 5.6, harmony 2.0, UCell 2.16, scDblFinder 1.26, NMF 0.28, clusterProfiler 4.20, edgeR 4.10, limma 3.68, BayesPrism 2.2, MuSiC 1.0, sandwich 3.1 and lmtest 0.9. Figures were generated using ggplot2 4.0 and ComplexHeatmap 2.28 ^25^. Exact package versions were recorded in the analysis environment lock file. Python 3.12.3 was used for file handling and table assembly.

### Data and code availability

All datasets analysed in this study are publicly available or access-controlled through the sources described above. MP gene lists, association results and summary statistics are provided in Supplementary Tables S1–S11. Donor-level data derived from HPAP are subject to the PANC-DB data-use terms, and access to controlled HPAP data is subject to the requirements of PANC-DB.

## Results

### Donor-resolved analysis identifies 43 recurrent transcriptional meta-programs across human islet cell types

To define recurrent transcriptional programs across human islet cell types, we analysed single-cell RNA-sequencing data from 48 donors in the discovery cohort GSE221156: 17 without diabetes (ND), 14 with prediabetes (PD) and 17 with type 2 diabetes (T2D). Of 245,878 donor-assigned cells, 208,976 (85.0%) passed quality control (Fig. S1A–C). Marker-based annotation assigned 200,403 cells to seven principal populations: α, β, δ, γ (pancreatic polypeptide), ductal, acinar and stellate cells (Fig. 1A, B; Fig. S2A). Each population showed the expected marker expression profile (Fig. S2B), and 91–100% of cells in each population carried the corresponding label in the original study (Fig. S2C). The remaining 8,573 cells were classified as uncertain or multihormonal and were not included in program discovery. In the original annotation, the multihormonal cells mapped predominantly to α-cell annotations, whereas the uncertain cells consisted mainly of endothelial, stellate and immune cells (Fig. S2B, C). Cell-type composition varied considerably between donors within each diagnostic group (Fig. S2D).

**Figure 1.**
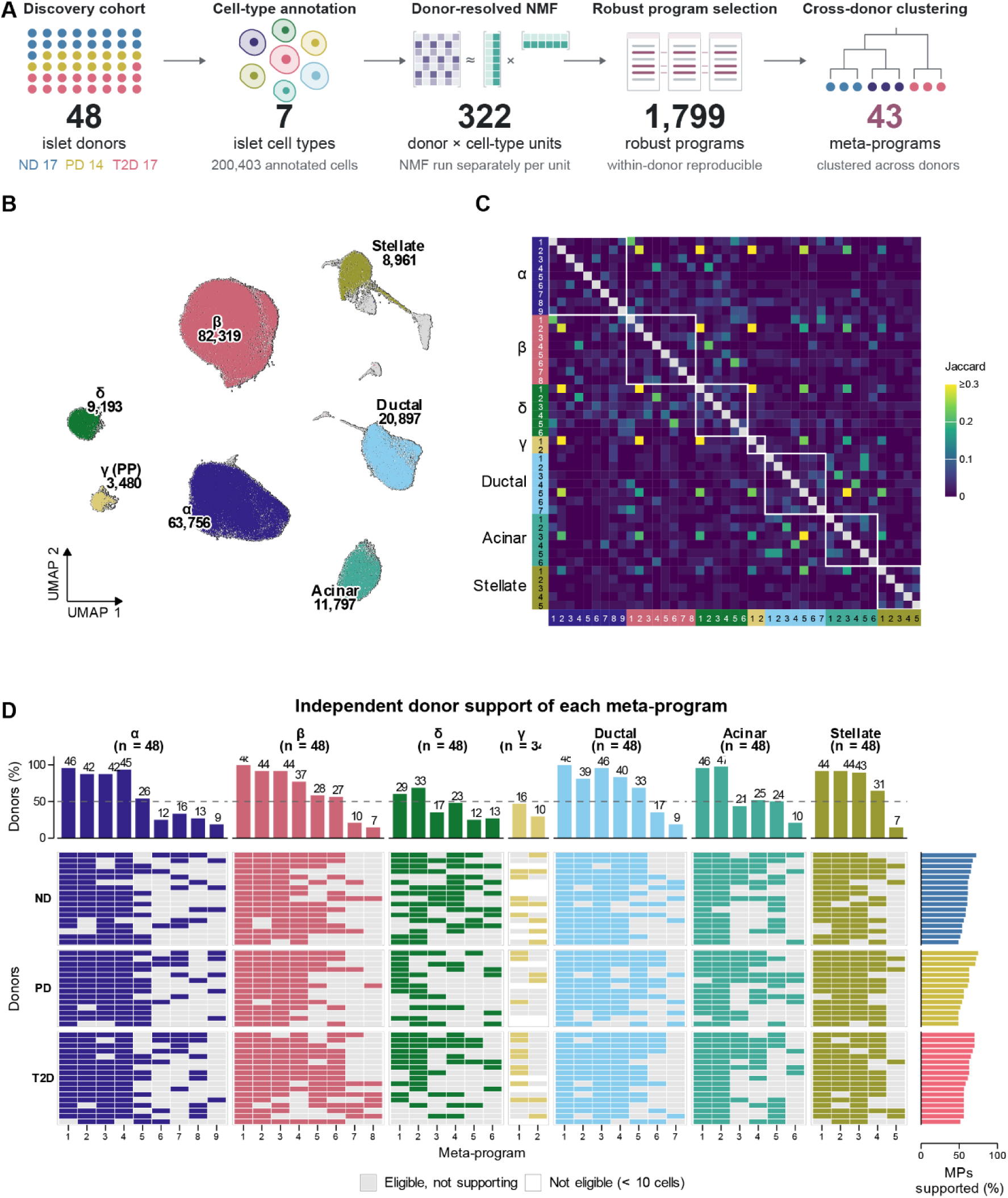
Donor-resolved meta-program analysis identifies recurrent transcriptional programs across human islet cell types. (A) Overview of the discovery workflow and key outputs from GSE221156, comprising 48 donors (17 ND, 14 PD and 17 T2D). NMF was performed separately for each donor × cell-type unit with at least 10 cells, yielding 1,799 robust programs that were clustered into 43 meta-programs (MPs), each represented by 50 genes. MP gene sets and scoring procedures were frozen before disease-association and validation analyses. (B) UMAP of 208,976 QC-passed cells coloured by cell type; cell numbers are indicated. Grey cells represent uncertain or multihormonal populations excluded from program discovery. (C) Pairwise gene overlap among the 43 MPs, measured by the Jaccard index. The hypoxia-response program shared across six cell types and the integrated stress-response program shared by α-MP1 and β-MP1 are indicated. (D) Donor support for each MP. Coloured tiles indicate donors contributing at least one robust program, grey tiles indicate eligible donors without a contributing program, and white tiles indicate donors with fewer than 10 cells of the corresponding cell type. Bars show the percentage of eligible donors supporting each MP. ND, donors without diabetes; PD, prediabetes; T2D, type 2 diabetes; NMF, non-negative matrix factorization.

We next applied non-negative matrix factorization (NMF) separately to each donor and cell type. The analysis covered 322 donor × cell-type units with at least 10 cells: all 48 donors for six cell types and 34 donors for γ cells (Fig. 1A; Fig. S3H). Across NMF ranks 4–9, these units yielded 12,558 candidate programs. Of these, 1,799 were classified as robust: they were reproducibly identified across NMF ranks within a donor, matched a program identified in at least one other donor and were non-redundant within that donor (Fig. S3I). The robust programs comprised 350 α-cell, 353 β-cell, 175 δ-cell, 71 γ-cell, 310 ductal, 278 acinar and 262 stellate programs. Within each cell type, these programs formed well-separated blocks of overlapping programs contributed by different donors (Fig. S3A–G).

Cross-donor clustering of the robust programs identified 43 meta-programs (MPs), each represented by 50 genes: nine α-cell, eight β-cell, six δ-cell, two γ-cell, seven ductal, six acinar and five stellate MPs (Fig. 1A, 1C; Fig. S3A–G, Table S1). Together, the MPs incorporated 1,465 robust programs, accounting for 81.4% of all robust programs (Fig. S3I, J). Each MP was supported by programs from 7–48 distinct donors, corresponding to 15–100% of the donors eligible for the respective cell type (Fig. 1D). Notably, 26 of the 43 MPs were supported by at least half of the eligible donors. This support was distributed broadly across the cohort: each donor contributed at least one robust program to a median of 61% of the MPs defined for its eligible cell types (range, 49–74%; Fig. 1D). Within each cell type, MPs were largely distinct, with a median pairwise gene overlap of one (Fig. 1C).

### β-cell meta-programs capture diverse functional features and donor-level expression patterns

To characterize the biological features of the islet meta-programs, we first tested all 43 MPs for enrichment of MSigDB Hallmark gene sets. Thirty-one MPs were significantly enriched for at least one gene set. Together, they represented diverse functional themes spanning metabolism, signaling, stress responses, immune processes, development and proliferation (Fig. 2A). Several themes recurred across cell types: epithelial– mesenchymal transition was enriched in 14 MPs, hypoxia and mTORC1 signaling in 12 each, TNF-α signaling via NF-κB in eight and glycolysis in seven. Full Hallmark, KEGG and GO:BP enrichment results are provided in Table S2, with GO:BP and KEGG enrichment profiles shown in Fig. S4 and S5.

**Figure 2.**
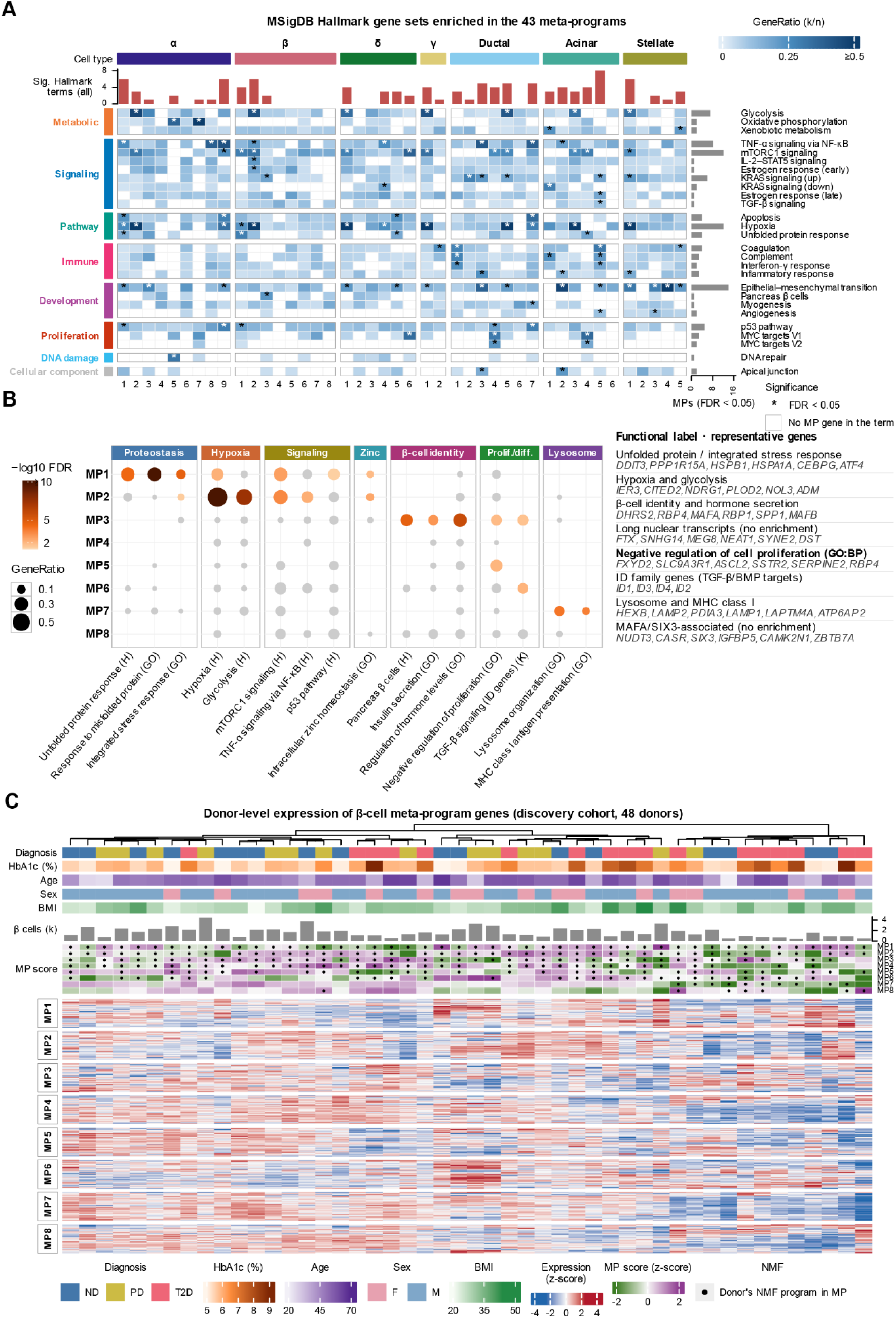
Functional characterization of islet meta-programs and donor-level expression of β-cell programs.(A) Hallmark gene-set enrichment across the 43 MPs. Rows represent gene sets significantly enriched in at least one MP (FDR < 0.05), grouped by biological category. Colour indicates GeneRatio, and asterisks indicate significant enrichment. (B) Functional annotation of the eight β-cell MPs based on enriched biological processes and representative member genes. Dot size and colour indicate enrichment strength. (C) Donor-level expression of the genes defining the eight β-cell MPs in the discovery cohort (48 donors). Columns represent donors, grouped by diagnosis; rows represent MP genes grouped by program. Expression is shown as gene-wise standardized pseudobulk expression. Clinical and donor-level annotations are displayed above the heatmap.

We next characterized the eight β-cell MPs in greater detail using Hallmark, KEGG and GO:BP enrichment analyses together with their member genes (Fig. 2B; Tables S2). Three β-cell MPs showed clear functional signatures. β-MP1 was enriched for the unfolded protein response, mTORC1 signaling and hypoxia. At the GO:BP level, it was further enriched for responses to misfolded proteins, protein refolding and cellular responses to unfolded proteins. Its representative genes included the integrated stress response-associated genes *DDIT3*, *PPP1R15A*, *CEBPG* and *ATF4*, together with the chaperones *HSPB1* and *HSPA1A*. β-MP2 showed particularly strong enrichment for hypoxia (18 of 28 annotated genes; FDR = 2.4 × 10⁻¹⁶) and glycolysis. It was also enriched for responses to oxygen levels, with representative genes including *IER3*, *CITED2*, *NDRG1*, *PLOD2*, *NOL3* and *ADM*. β-MP3 was enriched for the Hallmark pancreatic β-cell gene set, regulation of hormone levels and insulin secretion, and overlapped the KEGG maturity-onset diabetes of the young pathway. Representative β-MP3 genes included *DHRS2*, *RBP4*, *MAFA*, *RBP1*, *SPP1* and *MAFB*. The enrichment of the Hallmark pancreatic β-cell gene set was supported by six overlapping genes: *PDX1*, *PCSK2*, *G6PC2*, *MAFB*, *IAPP* and *ABCC8*.

β-MP6 and β-MP7 captured additional, more specific functional features. β-MP6 was enriched for KEGG TGF-β signaling, supported by the inhibitor-of-DNA-binding genes *ID1*–*ID4*, and for gap junction, driven by tubulin-family genes. β-MP7 showed enrichment for lysosomal function and antigen processing and presentation, supported by lysosomal genes (*HEXB*, *LAMP1*, *LAMP2*, *LAPTM4A* and *ATP6AP2*), together with *PDIA3* and the MHC class I genes *HLA-A*, *HLA-B* and *HLA-C*. β-MP4 and β-MP8 displayed distinct gene compositions that were not significantly represented by the tested gene-set collections. β-MP4 contained long noncoding and nuclear transcripts, including FTX, SNHG14, MEG8 and NEAT1, whereas β-MP8 included *CASR*, *SIX3*, *CAMK2N1* and *MAFA*. β-MP5 showed enrichment for negative regulation of cell population proliferation (GO:BP; 10 of 44 annotated genes; FDR = 0.009), supported by *FXYD2*, *SLC9A3R1*, *ASCL2* and *SSTR2*. Together, the eight β-cell MPs encompassed diverse gene compositions and functional features, including stress responses, hypoxia and glycolysis, β-cell identity, signaling and lysosomal processes.

We then examined the expression of all 400 genes comprising the eight β-cell MPs across the 48 donors in the discovery cohort (Fig. 2C). The heatmap revealed coordinated donor-level expression patterns among genes within individual MPs. Mean expression across the member genes of each MP correlated with its corresponding MP score (Pearson r = 0.89–0.98 for seven MPs and 0.67 for β-MP4). The number of donors contributing at least one robust NMF program also varied across MPs: all 48 donors contributed a program to β-MP1, compared with 10 donors for β-MP7 and seven for β-MP8. The predefined donor-level MP scores were subsequently used to quantify disease-associated variation.

### A β-cell meta-program is markedly reduced in type 2 diabetes

We next tested the 43 meta-programs for associations with diagnostic group and HbA1c in the discovery cohort, using donor-level pseudobulk scores from each program’s cell type of origin. Models were adjusted for age, sex and BMI. This screen identified two β-cell programs with significant T2D–ND differences: β-MP5, with lower scores in T2D, and β-MP3, with higher scores. None of the 35 programs from the other six cell types showed significant T2D–ND differences after FDR correction (Fig. 3A, B; Table S3).

**Figure 3.**
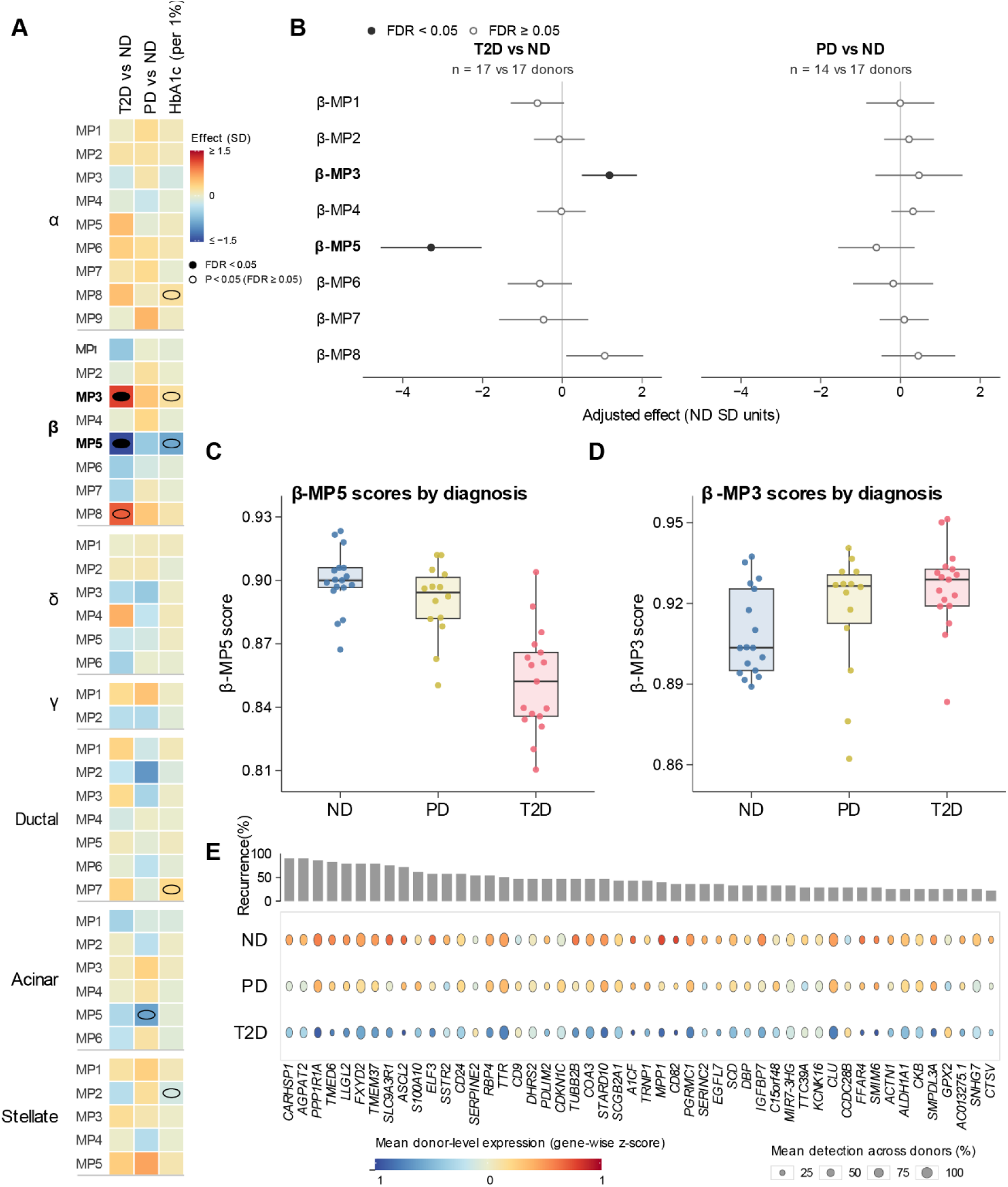
β-MP5 is reduced and β-MP3 is increased in T2D among 43 islet meta-programs. (A) Associations of all 43 MPs with T2D, prediabetes and HbA1c in the discovery cohort. Scores were calculated from donor-level pseudobulk expression using the frozen gene sets. Models were adjusted for age, sex and BMI. Effects are standardized to the ND donor score SD; the colour scale is truncated at ±1.5 SD. Solid circles indicate FDR < 0.05, and open circles indicate nominal P < 0.05 with FDR ≥ 0.05. The three association analyses represent different contrasts and are not directly comparable. (B) Adjusted effects of T2D versus ND and PD versus ND on the eight β-cell MP scores, expressed in ND donor SD units. Points indicate effect estimates and horizontal lines indicate 95% confidence intervals. (C, D) Donor-level β-MP5 and β-MP3 scores across diagnostic groups. Each point represents one donor; adjusted T2D-versus-ND effects are annotated. (E) Expression of the 50 β-MP5 member genes across ND, PD and T2D donors. Genes are ordered by their frequency among contributing NMF programs. Colour indicates mean gene-wise standardized donor-level expression, and dot size indicates the mean percentage of β cells expressing each gene. Group sizes: 17 ND, 14 PD and 17 T2D donors. Full estimates and statistical results for all 129 tests are provided in Table S3. HC3 robust standard errors and Benjamini–Hochberg FDR correction were used.

β-MP5 showed the largest absolute standardized T2D–ND effect among the 43 MPs. Scores were lower in T2D donors by 3.29 standard deviations of the ND donor distribution (adjusted effect, −3.29 SD; 95% CI, −4.56 to −2.03; FDR = 1.7 × 10⁻⁴; Fig. 3B, C). The reduction was consistent across individual donors: 16 of 17 T2D donors scored below the ND median, and 13 scored below the lowest ND value. The effect was robust to removal of any single donor: leave-one-donor-out analyses across the 34 ND and T2D donors yielded estimates ranging from −3.63 to −3.11 SD, with all 95% confidence intervals excluding zero (Fig. S6A). PD donors showed an intermediate score distribution, with a smaller adjusted difference from ND donors (−0.60 SD; FDR = 0.55).

β-MP3 changed in the opposite direction, with higher scores in T2D than in ND donors (adjusted effect, +1.19 SD; 95% CI, 0.49 to 1.88; FDR = 0.012; Fig. 3B, D). Sixteen of the 17 T2D donors scored above the ND median, and leave-one-donor-out estimates remained positive, ranging from +1.06 to +1.32 SD (Fig. S6B). These opposing changes in two β-cell transcriptional programs indicate that the disease-associated alterations were program-specific.

The HbA1c analysis showed a directionally consistent pattern. Across the 48 donors, higher HbA1c was associated with lower β-MP5 scores at nominal significance (adjusted effect, −0.89 SD per percentage-point increase in HbA1c; 95% CI, −1.47 to −0.32; P = 0.0031; FDR = 0.057; Fig. S7A, B). β-MP3 showed a positive association with HbA1c (+0.20 SD per percentage-point increase; P = 0.047; FDR = 0.41; Fig. S7A, C).

We further examined the expression of all 50 genes comprising β-MP5 across diagnostic groups (Fig. 3E). Forty-six of the 50 genes showed lower mean donor-level expression in T2D than in ND donors, indicating that the reduction extended across most members of the program rather than being confined to a small subset of genes. These included recurrent program members such as *CARHSP1*, *AGPAT2*, *PPP1R1A*, *FXYD2*, *TMEM37* and *SLC9A3R1*. Together, the donor-level score differences, their robustness to individual donors and the broad gene-level expression pattern identify β-MP5 as the principal T2D-associated β-cell transcriptional program in the discovery cohort.

### The T2D-associated reduction of β-MP5 is independently reproduced in human pancreatic β cells

To independently validate the β-cell meta-programs identified in the discovery cohort, we analysed single-cell RNA-sequencing data from the Human Pancreas Analysis Program (HPAP). The HPAP cohort comprised 58 donors (36 without diabetes and 22 with type 2 diabetes), contributing 40,876 β cells (Fig. 4A; Fig. S8A–C). To ensure adequate β-cell representation for donor-level scoring, the primary analysis included donors with at least 30 β cells, yielding 55 donors (34 ND and 21 T2D; Fig. S8D, E;). We applied the β-MP gene signatures to donor-level β-cell pseudobulk profiles. All 50 genes comprising β-MP5 were represented in the HPAP expression matrix (Fig. S8F; Table S4).

**Figure 4.**
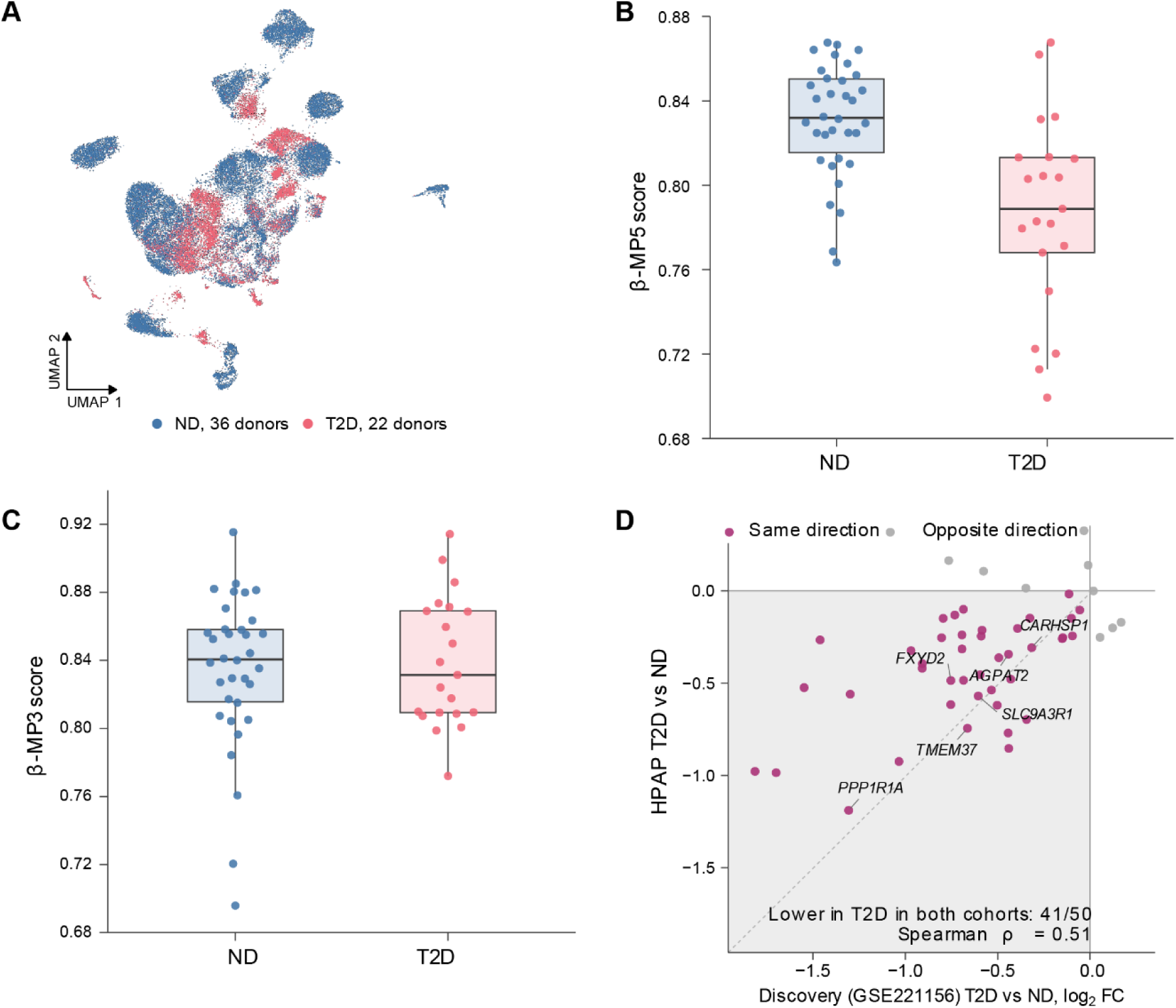
β-MP5 is reduced in T2D in an independent single-cell cohort (HPAP). (A) UMAP of 40,876 QC-passed β cells from 58 HPAP donors, coloured by diagnostic group (36 ND and 22 T2D). (B, C) Donor-level β-MP5 and β-MP3 scores in the primary validation analysis, including 55 donors with at least 30 profiled β cells (34 ND and 21 T2D). Adjusted effects, 95% confidence intervals, P values and FDR values are indicated. (D) Gene-level concordance of T2D-versus-ND expression changes for the 50 β-MP5 member genes between the discovery and HPAP cohorts. Each point represents one gene; purple indicates concordant directions and grey indicates opposite directions. Spearman correlations are shown for β-MP5 genes and, for comparison, all genes tested in both cohorts. Regression models were adjusted for age, sex, BMI and 10x chemistry. HPAP donors were independent of the discovery cohort.

β-MP5 scores were significantly lower in T2D donors in the HPAP cohort, reproducing the direction of the discovery result (adjusted effect, −1.68 SD of the ND donor distribution; 95% CI, −2.71 to −0.66; P = 0.0018; FDR = 0.015; Fig. 4B). Eighteen of the 21 T2D donors scored below the ND median.

In contrast, β-MP3 did not reproduce its discovery-stage increase in HPAP (adjusted effect, −0.36 SD; 95% CI, −0.90 to 0.17; P = 0.18; FDR = 0.48; Fig. 4C; Table S4). The two programs therefore showed distinct replication patterns, with β-MP5 emerging as the consistently supported T2D-associated program in the primary HPAP analysis.

We next examined whether the reduction in β-MP5 extended to its constituent genes across the two cohorts. Of the 50 β-MP5 genes, 46 had lower expression in T2D in the discovery cohort and 45 in HPAP, with 41 showing lower expression in both (Fig. 4D; Table S5). Gene-level T2D–ND log₂ fold changes were positively correlated between cohorts (Spearman ρ = 0.51), compared with a correlation of 0.16 across all genes tested in both cohorts. These findings extend the replication of β-MP5 from the program-score level to its constituent genes.

To further assess reproducibility, we analysed three additional human islet single-cell datasets generated using Smart-seq2: GSE153855, E-MTAB-5061 and GSE214517. Given their shared research origin and relatively small individual sample sizes, the datasets were pooled for validation (Fig. S9A–C; Table S6). Together, they comprised 27 donors, 7,254 quality-controlled cells and 1,103 β cells. Nineteen donors (11 ND and 8 T2D) had at least 30 β cells and were included in the pooled analysis. In a model adjusting for dataset, β-MP5 scores were significantly lower in T2D (adjusted effect, −2.03 SD; 95% CI, −2.87 to −1.18; P = 1.3 × 10⁻⁴; FDR = 0.001; Fig. S9D). The effect was consistently negative across the three datasets and remained robust in all three leave-one-dataset-out analyses, with estimates ranging from −2.18 to −1.90 SD and confidence intervals excluding zero.

β-MP3 scores were higher in T2D in the pooled Smart-seq2 analysis (adjusted effect, +1.36 SD; 95% CI, 0.36 to 2.36; P = 0.011; FDR = 0.022; Fig. S9E). However, this association contrasted with the HPAP result, indicating less consistent replication across independent cohorts. Taken together, the reduction of β-MP5 was reproduced in two independent single-cell validation settings, HPAP and the pooled Smart-seq2 datasets, and was accompanied by concordant changes across its constituent genes. These findings establish β-MP5 as a reproducible T2D-associated β-cell transcriptional program across independent human islet cohorts.

### Bulk islet transcriptomes reveal concordant reductions in the β-MP5 signature in type 2 diabetes

To extend the single-cell findings to bulk human islet transcriptomes, we analysed five publicly available datasets spanning microarray and RNA-sequencing platforms and different tissue preparations (Fig. 5A; Table S7). Three datasets profiled isolated islets, one profiled laser-capture microdissected (LCM) islets from living patients undergoing partial pancreatectomy, and one profiled LCM β-cell-enriched tissue. Four datasets provided ND and T2D comparisons, while GSE50244 provided HbA1c measurements for 77 donors. The 50-gene β-MP5 signature was applied across datasets, with gene coverage ranging from 38 to 50 members. GSE25724, which measured 38 genes (76%), was analysed only as a sensitivity estimate.

**Figure 5.**
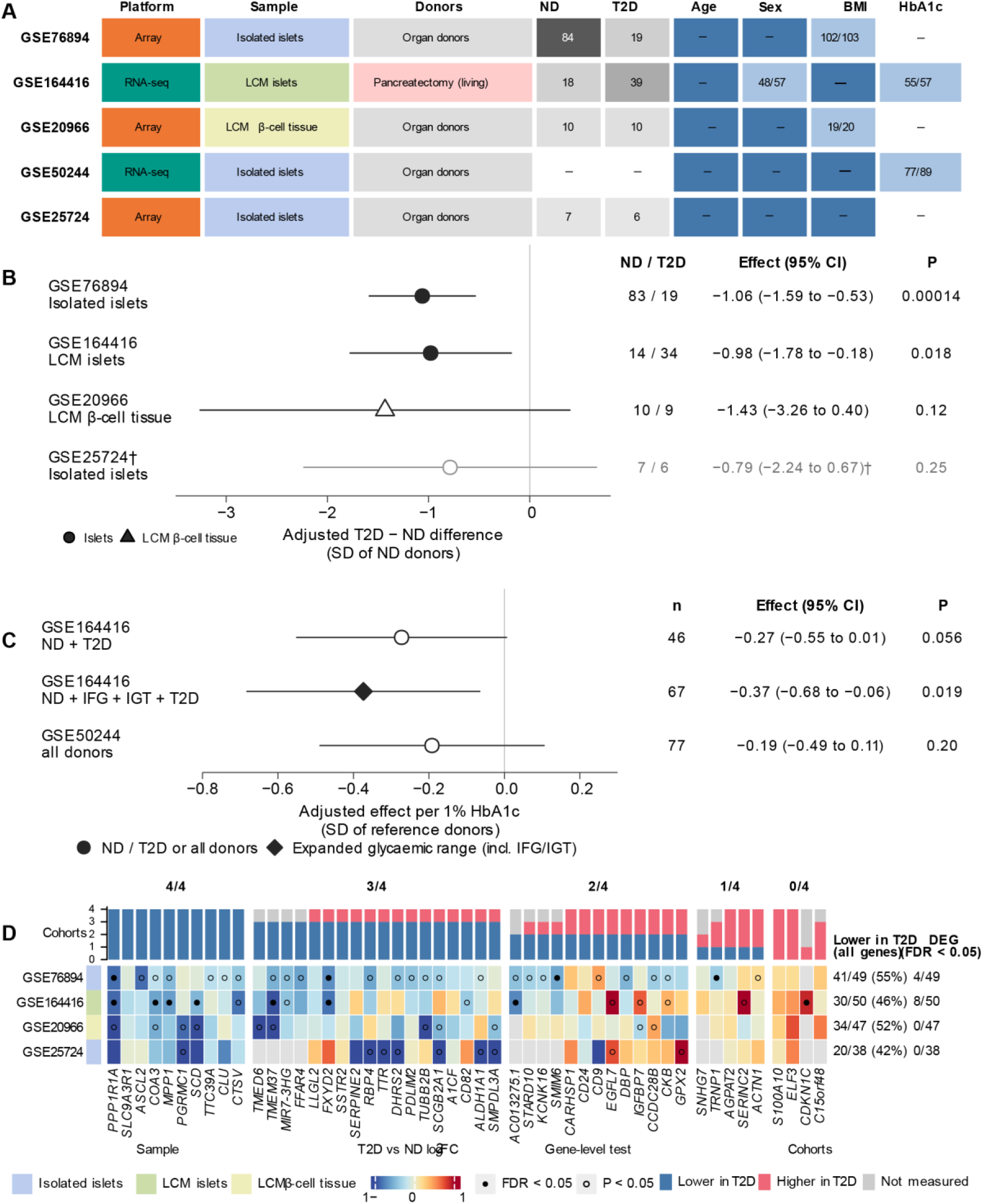
Bulk pancreatic tissue transcriptomes support the reduction of β-MP5 in T2D. (A) Characteristics of the five bulk transcriptomic cohorts, including platform, tissue preparation, donor source, diagnostic groups and availability of clinical variables. (B) Adjusted differences in β-MP5 scores between T2D and ND donors across four cohorts with diagnostic information. Effects are expressed in units of the corresponding ND donor score SD; points indicate adjusted estimates and horizontal lines indicate 95% confidence intervals. GSE25724, which measured 38 of the 50 β-MP5 genes, is shown as a sensitivity analysis. (C) Associations between β-MP5 scores and HbA1c in GSE164416 and GSE50244. The expanded glycaemic-range analysis in GSE164416 additionally included donors with impaired fasting glucose or impaired glucose tolerance. Effects represent changes per 1-percentage-point increase in HbA1c. (D) Gene-level expression changes of the 50 β-MP5 genes across the four diagnostic cohorts. Genes are grouped by the number of cohorts showing lower expression in T2D (4/4 to 0/4). Colour indicates T2D-versus-ND log₂ fold change; grey indicates genes not measured. Upper bars summarize directional changes across cohorts, and right-side annotations show the number and percentage of genes with lower expression in each cohort, together with the number of genome-wide significant differentially expressed genes. Models were adjusted for available donor covariates. Full results are provided in Supplementary Tables S7–S9.

β-MP5 scores were lower in T2D than in ND donors in all four datasets with diagnostic information (Fig. 5B). Significant reductions were observed in GSE76894 (adjusted effect, −1.06 SD; 95% CI, −1.59 to −0.53; P = 0.00014) and GSE164416 (−0.98 SD; 95% CI, −1.78 to −0.18; P = 0.018). The same direction was observed in GSE20966, comprising LCM β-cell-enriched tissue (−1.43 SD; 95% CI, −3.26 to 0.40; P = 0.12), and in the GSE25724 sensitivity analysis (−0.79 SD; 95% CI, −2.24 to 0.67; P = 0.25). Thus, the reduction of β-MP5 scores was directionally consistent across isolated islets, LCM islets and β-cell-enriched tissue, with the two larger cohorts providing statistically significant support.

We next examined the association between β-MP5 scores and HbA1c in the two datasets with available measurements (Fig. 5C). In GSE164416, higher HbA1c was associated with lower β-MP5 scores among ND and T2D donors (adjusted effect, −0.27 SD per percentage-point increase in HbA1c; 95% CI, −0.55 to 0.01; P = 0.056). In a prespecified sensitivity analysis that extended the population to donors with impaired fasting glucose (IFG) or impaired glucose tolerance (IGT), the inverse association was stronger (−0.37 SD per percentage-point increase; 95% CI, −0.68 to −0.06; P = 0.019). GSE50244 showed a concordant negative effect estimate (−0.19 SD per percentage-point increase; 95% CI, −0.49 to 0.11; P = 0.20). The inverse direction in both cohorts, together with the association observed across the broader glycaemic range in GSE164416, links β-MP5 expression with glycaemic variation in human islets.

Analysis of the 50 β-MP5 member genes revealed widespread T2D-associated expression changes across the bulk datasets (Fig. 5D; Table S8). Lower expression in T2D was observed for 41 of 49 genes in GSE76894 (84%), 30 of 50 in GSE164416 (60%), 34 of 47 in GSE20966 (72%) and 20 of 38 in GSE25724 (53%). These proportions exceeded the corresponding proportions of genes with lower expression across the tested transcriptomes (55%, 46%, 52% and 42%, respectively). Ten β-MP5 genes, including *PPP1R1A*, *SLC9A3R1*, *SCD* and *CLU*, showed lower expression in T2D across all four datasets. Few member genes, however, showed FDR-significant differential expression on their own (FDR < 0.05 across all tested genes: 4 of 49 in GSE76894, 8 of 50 in GSE164416 and none in GSE20966 or GSE25724). The bulk-level reduction of β-MP5 therefore reflected coordinated, modest decreases across many constituent genes rather than large changes in a few genes that would be captured by conventional differential expression analysis.

We also assessed islet cell composition using reference-based deconvolution with BayesPrism, MuSiC and CIBERSORTx (Table S9). Estimated β-cell fractions were most interpretable in GSE164416, but their differences between diagnostic groups varied across methods, and comparable estimates were not available across all datasets. Accordingly, the primary bulk analyses used cohort-level expression models without estimated cell-fraction covariates.

Together, these analyses demonstrate consistent directional reductions of β-MP5 across multiple human islet bulk transcriptomic datasets and tissue preparations, with accompanying gene-level expression changes and inverse effect estimates for HbA1c. The bulk findings extend the single-cell observations to independent transcriptomic platforms and provide additional support for β-MP5 as a T2D-associated human islet transcriptional signature.

### β-MP5 tracks ex vivo insulin secretory function in human islets

To examine whether β-MP5 is related to β-cell function, we matched HPAP β-cell transcriptomes with islet perifusion measurements from the same donors (Fig. 6A; Table S10). Islets from HPAP donors were perifused in two laboratories: 52 donors at the University of Pennsylvania (UPenn; 32 ND and 20 T2D) and 42 donors at Vanderbilt University (27 ND and 15 T2D). Thirty-nine donors were measured in both laboratories, providing two laboratory measurements from largely overlapping donors. For each donor with at least 30 profiled β cells, we related the frozen β-MP5 score derived from the β-cell pseudobulk to insulin area under the curve (AUC) during stimulation with 16.7 mM glucose, glucose plus IBMX and KCl. Models were adjusted for age, sex, BMI and 10x chemistry.

**Figure 6.**
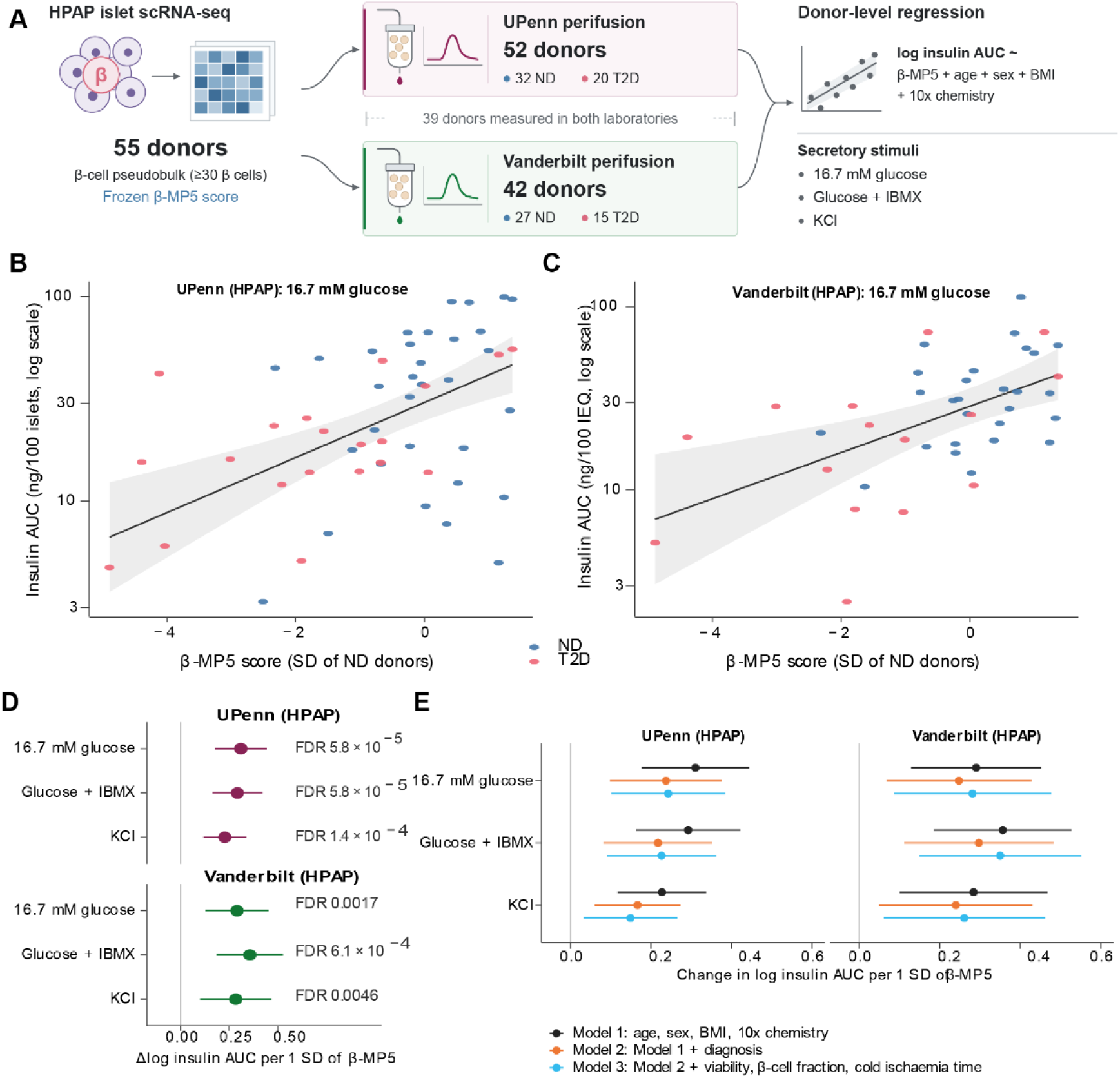
β-MP5 tracks ex vivo insulin secretory function in human islets. (A) Study design linking HPAP β-cell transcriptomes with donor-matched islet perifusion measurements from UPenn (52 donors) and Vanderbilt (42 donors). Thirty-nine donors had measurements from both laboratories; the two datasets therefore do not represent independent donor cohorts. (B, C) Associations between β-MP5 scores and insulin AUC during 16.7 mM glucose stimulation in UPenn and Vanderbilt, respectively. Each point represents one donor. Lines and shaded areas show model-adjusted estimates and 95% confidence bands. (D) Adjusted associations between β-MP5 scores and insulin AUC under glucose, glucose plus IBMX and KCl stimulation in both laboratories. Effects are expressed as changes in log insulin AUC per 1-SD increase in β-MP5 score, with 95% confidence intervals. (E) Associations under three nested models: Model 1, adjusted for age, sex, BMI and 10x chemistry; Model 2, additionally adjusted for diagnosis; and Model 3, additionally adjusted for islet viability, β-cell fraction and cold ischaemia time. Filled symbols indicate FDR < 0.05 within the corresponding testing family. All analyses were performed at the donor level using HC3 robust standard errors.

Higher β-MP5 scores were associated with greater glucose-stimulated insulin secretion in both laboratories (Fig. 6B, C). In UPenn, each 1-SD increase in β-MP5 corresponded to a 0.31 increase in log insulin AUC (95% CI, 0.18 to 0.44; FDR = 5.8 × 10⁻⁵). The association was closely replicated in Vanderbilt, with an adjusted effect of 0.29 (95% CI, 0.13 to 0.45; FDR = 0.0017). The positive association extended to glucose plus IBMX (UPenn, 0.29; Vanderbilt, 0.36) and KCl-induced depolarization (UPenn, 0.23; Vanderbilt, 0.28), with FDR ≤ 0.005 for all six associations (Fig. 6D; Fig. S10A–D). Thus, β-MP5 tracked insulin release across multiple secretory stimuli in measurements from both laboratories.

We next examined whether these associations were attributable to diagnostic group or islet preparation characteristics (Fig. 6E). Additional adjustment for diagnosis reduced the effect estimates modestly, but all remained positive, ranging from 0.17 to 0.24 in UPenn and from 0.24 to 0.30 in Vanderbilt. Further adjustment for islet viability, β-cell fraction and cold ischaemia time produced similar estimates (0.15–0.24 in UPenn and 0.26–0.35 in Vanderbilt; Table S11). These results indicate that the association between β-MP5 and insulin secretion persisted after accounting for diagnostic status and measured islet preparation characteristics. The associations were also robust to individual donors: leave-one-donor-out estimates ranged from 0.20 to 0.35 in UPenn and from 0.25 to 0.41 in Vanderbilt, with no donor having a Cook’s distance above 0.6 (Table S11).

We further examined whether β-MP5 was associated with other measures of β-cell function (Fig. S11; Table S11). Insulin content was not significantly associated with β-MP5 in Vanderbilt (adjusted effect, 0.09; 95% CI, −0.11 to 0.28; FDR = 0.38) and could not be estimated in UPenn, where only 16 donors had measurements. In contrast, the glucose stimulation index and the PANC-DB glucose-stimulated insulin response (GSIR) index showed positive associations with β-MP5 in the basic adjustment model (FDR = 0.0022 and 0.015, respectively). These associations were attenuated after adjustment for diagnosis (FDR = 0.19 and 0.096, respectively). Serum C-peptide also showed a positive association in the basic model (adjusted effect, 0.23; FDR = 0.015), which was markedly attenuated after diagnosis adjustment (adjusted effect, 0.03; FDR = 0.77). Together, these findings indicate that β-MP5 was more closely associated with stimulated insulin release than with insulin content. The association was also selective among the eight β-cell programs: none of the other seven MPs was associated with any perifusion AUC at FDR < 0.1 in either laboratory (smallest FDR = 0.14; Table S11).

To place β-MP5 in its tissue context, we mapped the 50-gene signature onto 16 spatial transcriptomic sections of whole pancreas from three NGT and three T2D donors (GSE264331; Fig. S12). β-MP5 scores varied across each section rather than being uniformly distributed (Fig. S12A). Islet-rich spots were identified from seven non-hormone endocrine marker genes, none of which is a β-MP5 member. In donor-level pseudobulk profiles, β-MP5 scores were higher in islet-rich than in endocrine-low spots in all six donors (exact sign-flip P = 0.031; Fig. S12B), indicating that β-MP5 expression in intact pancreatic tissue is concentrated in islet regions.

Collectively, these results connect β-MP5 with the secretory capacity of human β cells. Higher β-MP5 expression was associated with greater insulin release across multiple stimuli, with closely similar glucose-stimulated associations in two laboratories and persistence after adjustment for diagnosis and islet preparation characteristics. Together with its reproducible reduction in T2D, these findings identify β-MP5 as a transcriptional program associated with human β-cell secretory function.

## Discussion

In this study, we used donor-resolved NMF to define 43 recurrent transcriptional programs across seven human islet cell types and identified one β-cell program, β-MP5, that was markedly reduced in T2D. This reduction was reproduced in an independent single-cell cohort, in pooled Smart-seq2 datasets and, in the same direction, across four bulk islet transcriptomic datasets generated using different platforms and tissue preparations. Beyond its association with diagnosis, β-MP5 was associated with insulin secretion measured ex vivo in two laboratories. Higher scores were associated with greater insulin release in response to glucose, glucose plus IBMX and KCl, and these associations persisted after adjustment for diagnosis and islet preparation characteristics. None of the other seven β-cell programs was associated with perifusion AUC at FDR < 0.1. Together, these findings identify β-MP5 as a reproducible T2D-associated transcriptional program linked to stimulated insulin secretory capacity in human islets.

A central difficulty in islet transcriptomics has been the limited reproducibility of T2D-associated gene expression changes. Independent single-cell studies of human islets have reported largely non-overlapping sets of differentially expressed β-cell genes ^28–30^. Part of this discordance arises from analyses that treat cells as independent observations, which inflates the apparent precision of disease effects when the number of donors is small ^32,33^. Our design addressed this problem at two levels. First, programs were discovered within each donor and retained only when they recurred across donors, preventing a single donor with many cells from defining a program. Second, all disease and functional associations were tested using donor-level pseudobulk scores, with donors as the statistical unit, and with gene sets and a scoring procedure fixed before validation. The β-MP5 findings also illustrate the value of program-level analysis. Across the bulk cohorts, most β-MP5 genes showed lower expression in T2D and the program score was consistently reduced, although relatively few genes reached genome-wide significance individually. A consistent directional shift across many members of a fixed gene set can therefore be captured at the program level even when individual genes do not pass genome-wide thresholds, a property also exploited by meta-program analyses in cancer ^46^.

The reproducibility of β-MP5 across settings is notable because the validation data differed from the discovery cohort in several technical and biological respects. The single-cell validation cohorts used droplet-based and plate-based chemistries, while the bulk datasets included microarray and RNA-sequencing profiles of isolated islets from organ donors, laser-capture microdissected islets from living donors undergoing pancreatectomy and β-cell-enriched tissue from pancreas sections ^15,17,18,23,38^. Islets from living donors are not exposed to brain death, organ procurement or islet isolation, which are recognized sources of variability in donor islet studies ^20^. The directionally lower β-MP5 scores in these preparations suggest that the observed reduction is not explained solely by factors associated with organ procurement and islet isolation. The inverse HbA1c effect estimates in both bulk cohorts with available measurements, together with the significant association in GSE164416 when donors with impaired fasting glucose or impaired glucose tolerance were included, further suggest that β-MP5 varies along the glycaemic continuum rather than only between diagnostic categories.

The composition of β-MP5 points to features of a differentiated β-cell state associated with secretory function. Several of its members have established roles in insulin secretion. PPP1R1A is a MafA target gene that supports incretin-mediated amplification of glucose-stimulated insulin secretion, and its expression is reduced in islets from donors with T2D ^70^. PPP1R1A and TMEM37 were also among the 19 signature genes of T2D islets defined by the IMIDIA consortium, in which both genes modulated insulin secretion in β cells ^17^. STARD10 lies at a T2D risk locus, and reduced STARD10 expression is associated with defective insulin secretion in humans and mice ^71^. KCNK16 encodes the T2D-associated two-pore domain potassium channel TALK-1, which regulates β-cell electrical excitability and second-phase insulin secretion ^72^. Transthyretin (TTR) promotes glucose-induced calcium influx and insulin release in β cells ^73^, while FXYD2 has been identified as a β-cell-enriched component of the Na⁺/K⁺-ATPase ^74^.

CD9, another member, is one of the two surface markers that define human β-cell subtypes ^39^. β-MP5 also contains the cell-cycle inhibitor CDKN1C ^75^ and was enriched for genes involved in negative regulation of cell population proliferation. Together, these features are consistent with β-MP5 capturing aspects of a differentiated β-cell state associated with stimulus–secretion coupling. Its reduction in T2D is in line with models in which β cells lose aspects of their differentiated identity under metabolic stress ^8–10,41^, although our data do not establish which upstream factors drive this change.

The functional analyses provide the most direct evidence linking β-MP5 to β-cell performance. Similar positive associations were observed in perifusion measurements from two laboratories, despite differences in their experimental protocols and measurement units. The association extended from glucose to glucose plus IBMX, which amplifies secretion through cAMP signalling, and to KCl, which depolarizes the β-cell membrane and triggers insulin release largely independently of glucose metabolism ^76^. This pattern suggests that the association extends beyond glucose sensing alone and involves broader aspects of the stimulus–secretion pathway. In contrast, β-MP5 was not associated with islet insulin content, indicating that it relates more closely to stimulated insulin release than to the amount of stored insulin. The associations persisted after adjustment for diagnosis, islet viability, β-cell fraction and cold ischaemia time, indicating that they were not explained solely by diagnostic group differences or by the measured characteristics of the islet preparations, whose secretory profiles can vary widely ^19^. These findings extend previous efforts to connect islet transcriptomes with function, including Patch-seq studies relating single-cell gene expression to exocytosis ^48^ and multiomic studies of β-cell subtypes with distinct functional properties ^36^, by showing that a donor-level transcriptional program score is associated with whole-islet secretory output. Because none of the other β-cell MPs showed comparable associations, the relationship with secretory function was specific to β-MP5 among the recurrent β-cell programs rather than a general feature of them. By contrast, β-MP3, the second T2D-associated program in the discovery cohort, showed inconsistent replication: it was not increased in T2D in HPAP, although it was higher in T2D in the pooled Smart-seq2 datasets. This difference highlights the importance of independent validation across cohorts before assigning functional significance to disease-associated programs.

The spatial transcriptomic analysis added a tissue-level perspective. In pancreas sections from six donors, β-MP5 scores were higher in islet-rich spots than in surrounding endocrine-low tissue in every donor. Islet-rich spots were defined using non-hormone endocrine genes that are not members of β-MP5, reducing the potential for circularity and limiting the influence of the hormone transcript bleed reported for this dataset ^49^. Because spatial transcriptomic spots capture several cells ^77^, this analysis shows that β-MP5 expression is enriched in islet regions of intact pancreas but does not resolve its expression in individual β cells.

These findings have several translational implications. First, β-MP5 provides a compact, fixed gene set that can be scored in single-cell, bulk and spatial transcriptomic data without re-deriving the signature, making it a potentially useful common readout of β-cell-associated functional state across studies. Second, variability in human islet preparations is a recognized obstacle in islet research and transplantation ^19,20^. A transcriptional readout associated with secretory capacity could therefore complement existing functional assays when assessing islet preparation quality. Prospective studies will be needed to determine whether β-MP5 can improve the evaluation of islet preparations or predict functional outcomes after transplantation. Third, because β-cell function can partly recover in early T2D ^14^, β-MP5 may provide a molecular endpoint for experimental studies aimed at preserving or restoring β-cell function. Its constituent genes, including PPP1R1A, STARD10 and KCNK16, also provide candidates for future perturbation studies investigating the molecular basis of β-cell secretory dysfunction.

Several limitations should be considered. All human data were cross-sectional, so the analyses cannot determine whether reduced β-MP5 expression precedes, accompanies or results from β-cell dysfunction and hyperglycaemia. Most islets were obtained from organ donors, and residual confounding by donor characteristics, medication, terminal illness and procurement cannot be excluded. The two perifusion datasets measured largely overlapping donors and therefore represent two laboratory measurements rather than two independent donor cohorts. The discovery and validation cohorts were modest in size, and the pooled Smart-seq2 datasets came from a single research group. The spatial analysis was based on six donors and was therefore supportive rather than definitive. Some bulk cohorts measured only part of the β-MP5 gene set, and estimated cell-type composition differed across deconvolution methods ^28,68,69^. Consequently, the bulk associations could not be fully separated from differences in islet cellular composition. Because NMF was applied within each donor, the discovery step identifies programs that vary between cells of a donor and may miss changes that affect all β cells of a donor uniformly. T2D is also clinically heterogeneous ^78^, and we could not examine whether β-MP5 differs between clinical subgroups. Finally, our analyses establish associations; experimental perturbation of β-MP5 member genes in human β cells will be needed to determine whether components of this program contribute causally to secretory capacity.

In summary, donor-resolved analysis of human islet transcriptomes identified β-MP5 as a β-cell program that is reproducibly reduced in T2D across single-cell and bulk datasets, associated with ex vivo insulin secretion, and enriched in islet regions of intact human pancreas. β-MP5 provides a reproducible molecular readout associated with β-cell functional state and a starting point for mechanistic and translational studies of β-cell failure in T2D.

## Conclusion

Donor-resolved meta-program analysis of human islet transcriptomes identified β-MP5, a 50-gene transcriptional program derived from β cells that is reduced in T2D. This reduction was reproduced in independent single-cell datasets and was directionally consistent across bulk transcriptomic cohorts from organ donors and living donors.

Spatial transcriptomic analysis further showed that β-MP5 expression was enriched in islet regions of intact human pancreas. In donor-matched islets, higher β-MP5 scores were associated with greater insulin secretion in response to glucose, cAMP amplification and membrane depolarization, and these associations persisted after adjustment for diagnosis and measured islet preparation characteristics. Together, these findings establish β-MP5 as a reproducible transcriptional correlate of human β-cell secretory capacity. Its fixed gene set may provide a transferable molecular readout for studying β-cell functional states across cohorts and platforms. Such a readout could complement existing approaches to islet quality assessment and provide a potential molecular endpoint for studies aimed at preserving or restoring β-cell function in T2D. Longitudinal and perturbation studies will be needed to determine whether changes in β-MP5 contribute causally to β-cell dysfunction.

---

**Supplementary Figure 1.**
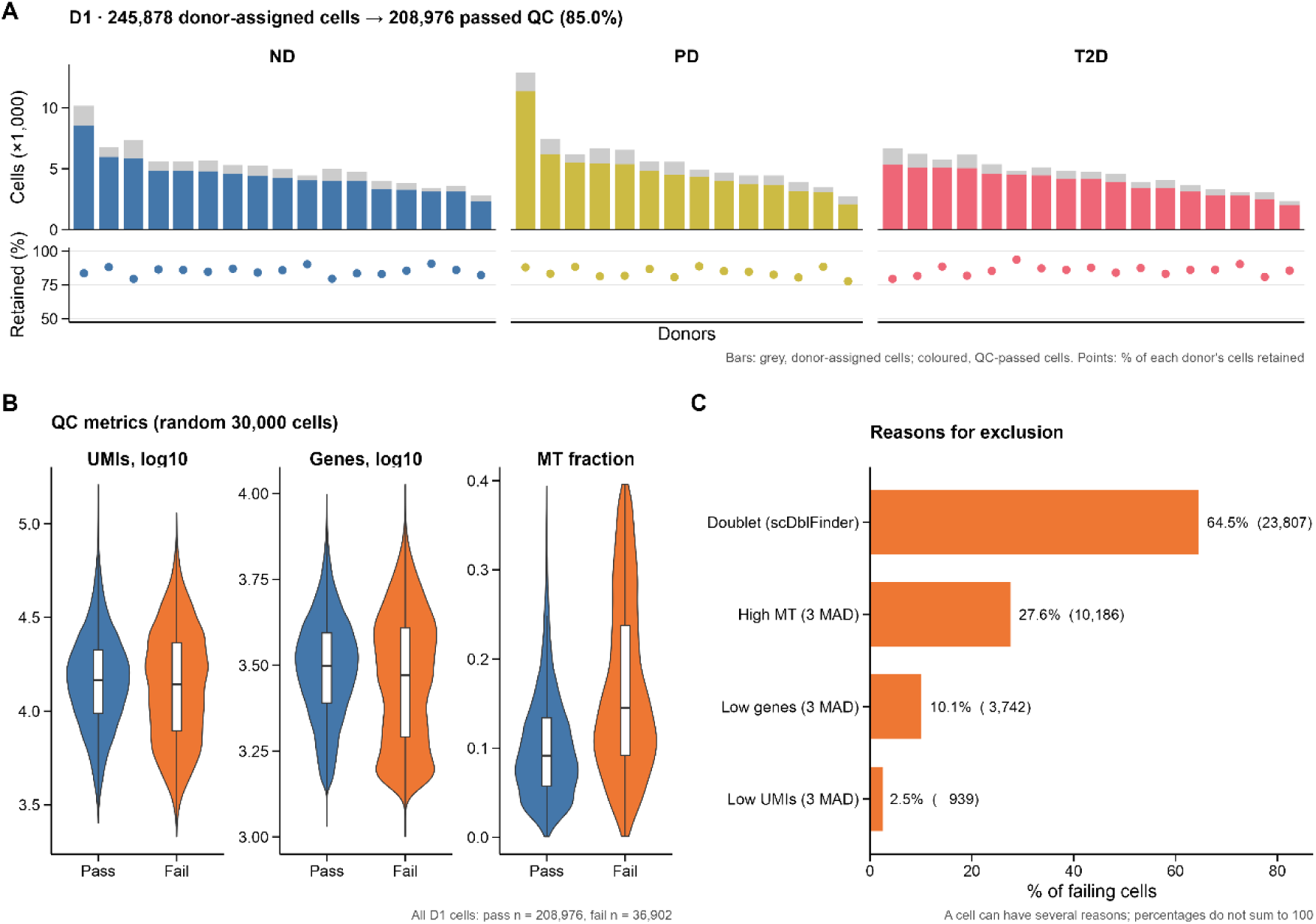
Quality control of the discovery cohort.(A) Number of cells per donor before and after QC. Grey indicates donor-assigned cells and colours indicate QC-passed cells; points show the percentage retained. (B) UMI counts, detected genes and mitochondrial transcript fractions in passing and failing cells (random sample of 30,000 cells). (C) Reasons for cell exclusion among the 36,902 failing cells. Cells could fail multiple criteria; percentages therefore do not sum to 100. Overall, 208,976 of 245,878 donor-assigned cells (85.0%) passed QC.

**Supplementary Figure 2.**
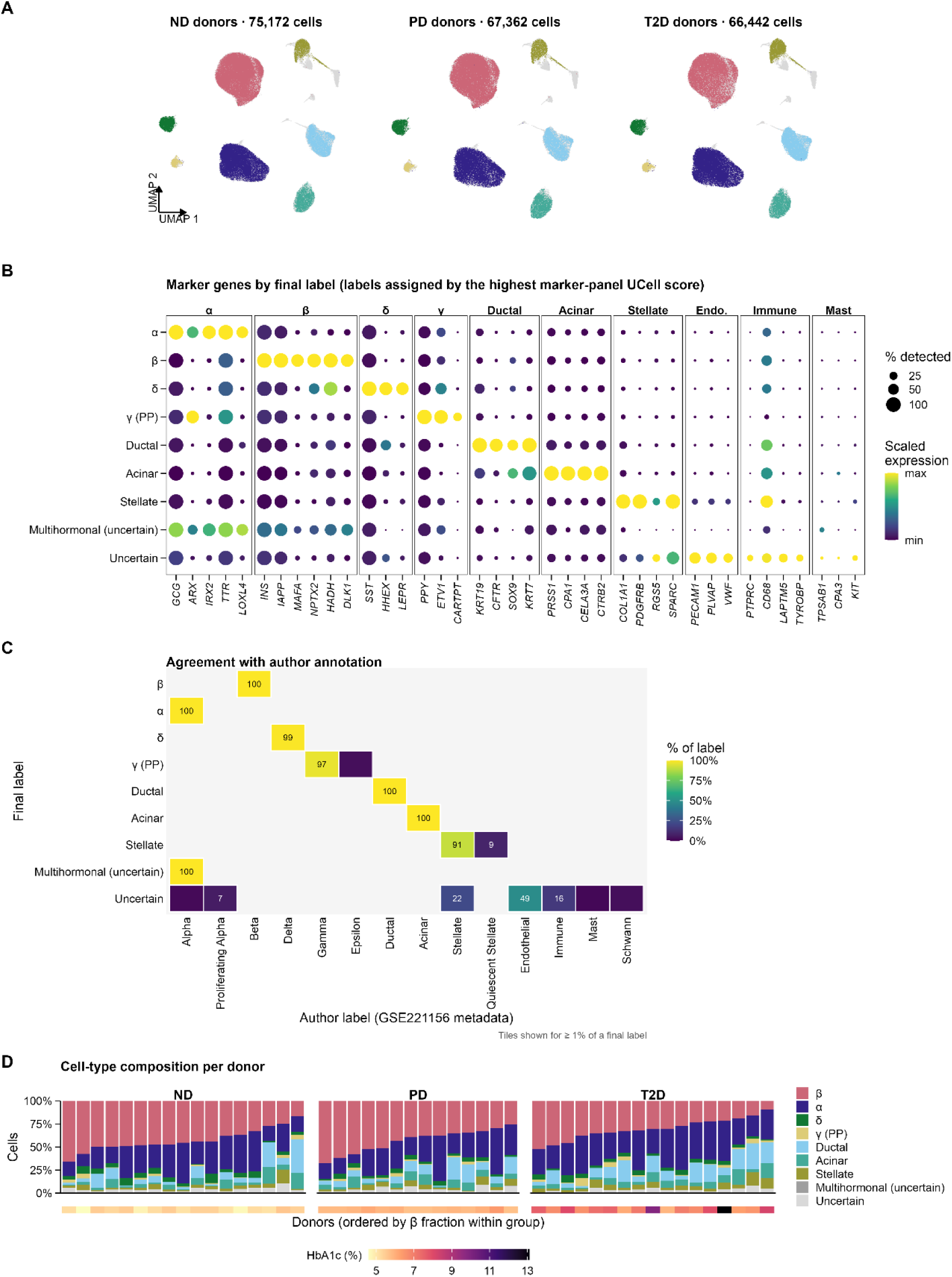
Cell-type annotation and cellular composition of the discovery cohort. (A) UMAPs showing cells from ND, PD and T2D donors separately. (B) Expression of marker-panel genes across the final cell-type labels. Dot size indicates the percentage of cells expressing each gene, and colour indicates scaled mean expression. Endothelial, immune and mast-cell panels are included to characterize uncertain cells excluded from program discovery. (C) Agreement between the final annotations and the original study annotations. (D) Cell-type composition of individual donors, ordered by β-cell fraction within each diagnostic group, with donor HbA1c shown below.

**Supplementary Figure 3.**
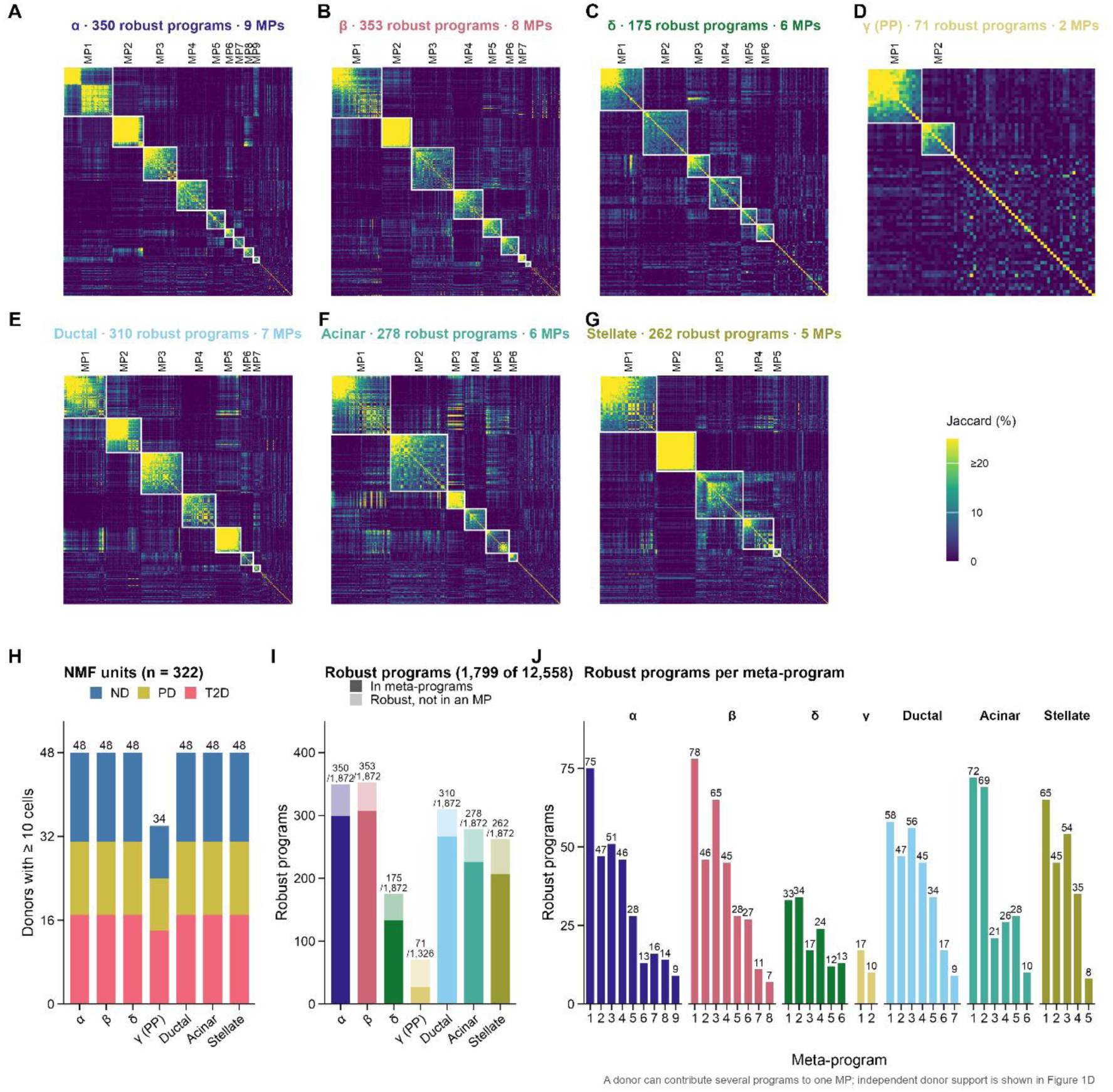
Construction and donor support of the 43 meta-programs. (A– G) Pairwise gene overlap among robust NMF programs within each cell type, measured by the Jaccard index. White boxes indicate programs assigned to the corresponding MPs. (H) Donor × cell-type units included in NMF (322 units). (I) Number of robust programs per cell type, divided into programs assigned and not assigned to an MP. A program was considered robust if it was reproduced at another NMF rank within the same donor (at least 35 of 50 genes shared), recurred in another donor (at least 10 genes shared) and was not redundant with a previously selected program from the same donor (at most 10 genes shared). (J) Number of robust programs contributing to each MP. A donor could contribute multiple programs to the same MP..

**Supplementary Figure 4.**
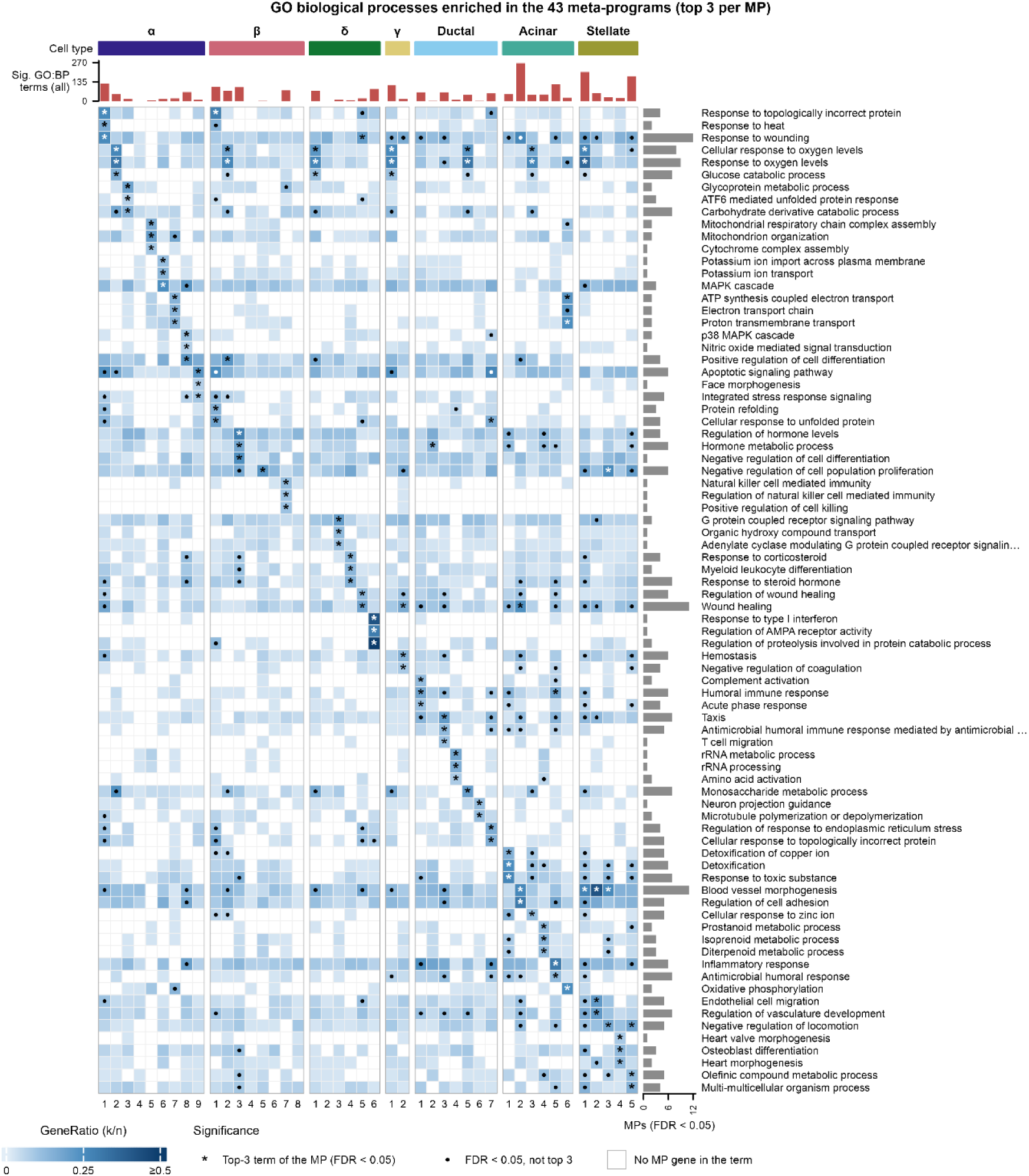
GO biological process enrichment across the 43 meta-programs.The union of the three most significant GO terms (FDR < 0.05) for each MP is shown. The 80 terms are ordered by the MP in which each term first appears among the top three enriched terms.

**Supplementary Figure 5.**
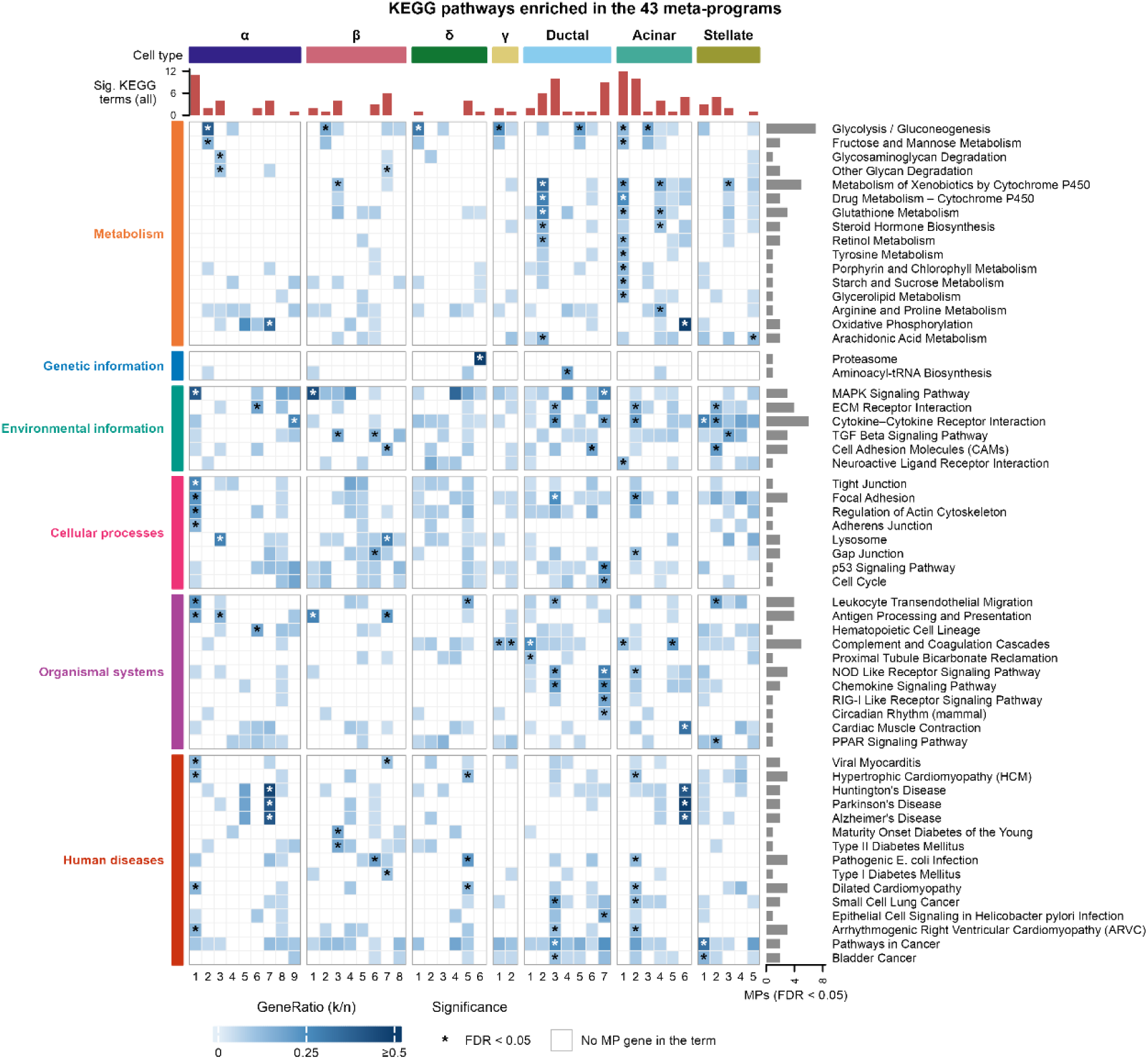
KEGG pathway enrichment across the 43 meta-programs. All 58 KEGG pathways significantly enriched in at least one MP (FDR < 0.05) are shown, grouped by KEGG top-level category. Asterisks indicate significant enrichment.

**Supplementary Figure 6.**
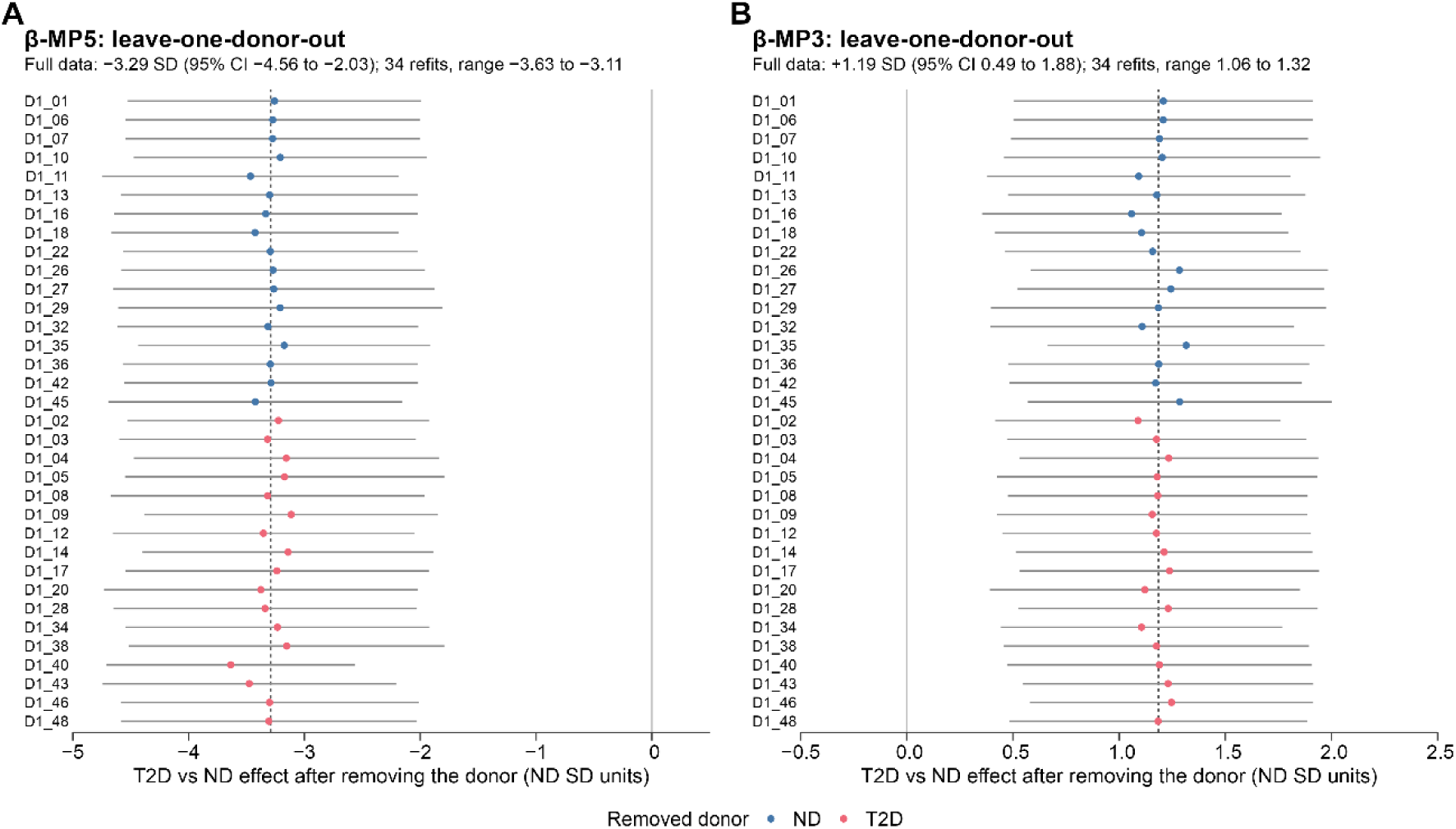
Sensitivity of β-MP5 and β-MP3 disease associations to individual donors. Leave-one-donor-out analyses of the adjusted T2D-versus-ND effects for β-MP5 (A) and β-MP3 (B). Each point represents an estimate obtained after removing one donor and refitting the regression model.

**Supplementary Figure 7.**
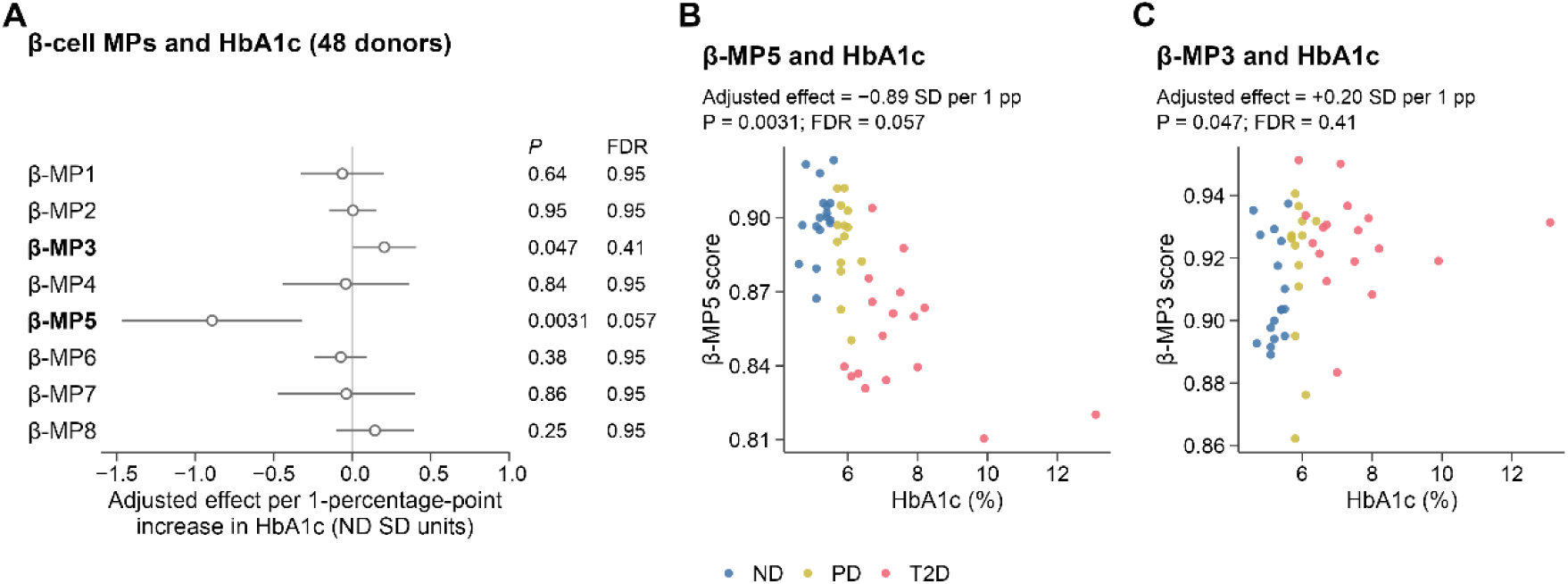
Associations of β-cell meta-program scores with HbA1c in the discovery cohort. (A) Adjusted effects of HbA1c on the eight β-cell MP scores, expressed per 1-percentage-point increase in HbA1c and standardized to the ND donor score SD. Points indicate effect estimates and horizontal lines indicate 95% confidence intervals. (B, C) Donor-level β-MP5 and β-MP3 scores plotted against HbA1c, with donors coloured by diagnostic group. Models were adjusted for age, sex and BMI.

**Supplementary Figure 8.**
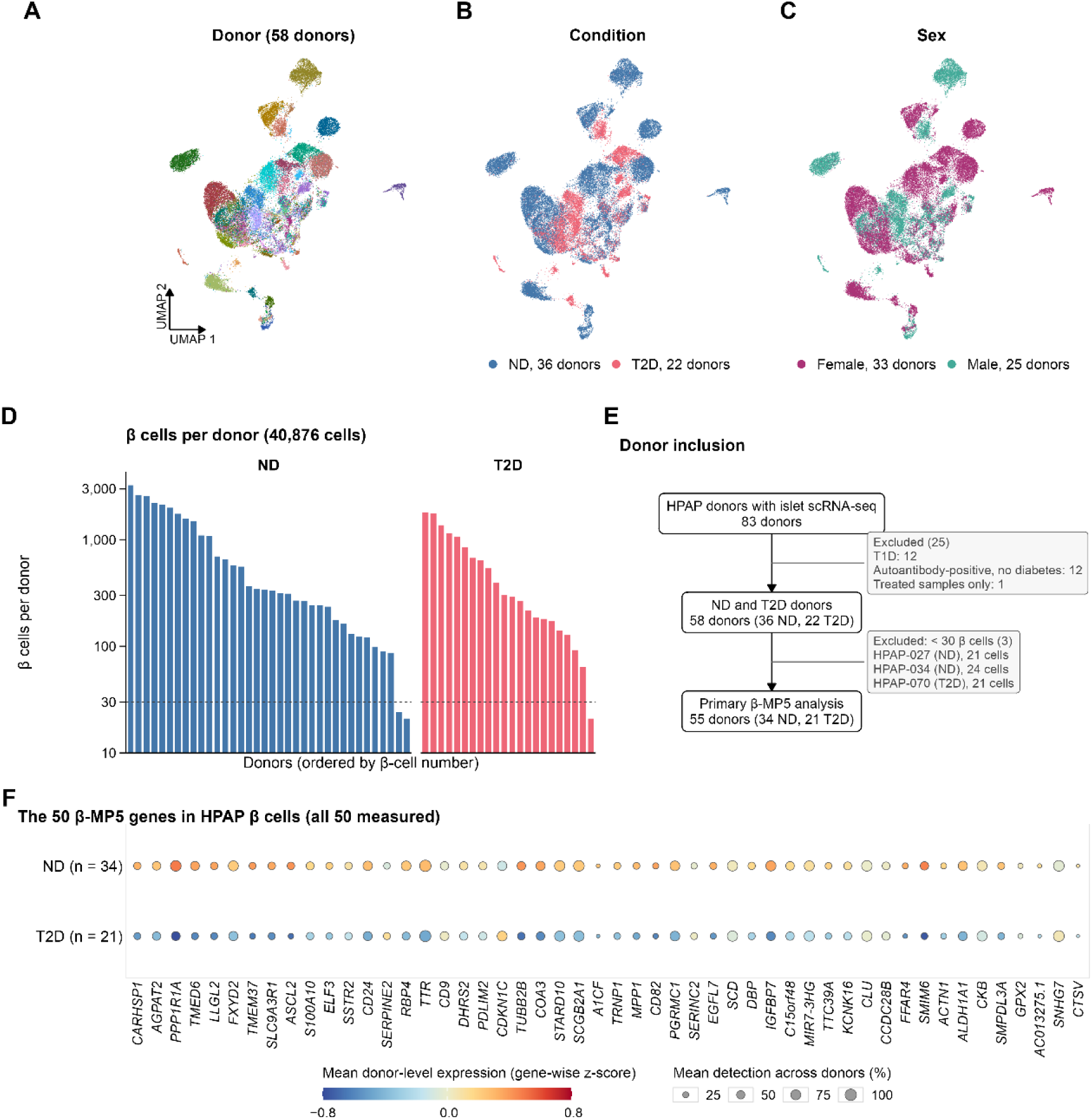
Quality control and donor composition of the HPAP β-cell validation cohort. (A–C) UMAP of 40,876 QC-passed β cells from 58 HPAP donors, coloured by donor (A), diagnosis (B) and sex (C). (D) Number of β cells per donor; the dashed line indicates the 30-cell threshold used in the primary validation analysis. (E) Donor inclusion and exclusion from the original 83 HPAP donors with islet scRNA-seq data. The primary analysis included 55 donors (34 ND and 21 T2D). (F) Expression of the 50 β-MP5 genes in the primary analysis cohort. Colour indicates mean donor-level standardized expression, and dot size indicates the mean percentage of β cells expressing each gene. Genes are ordered by their β-MP5 rank.

**Supplementary Figure 9.**
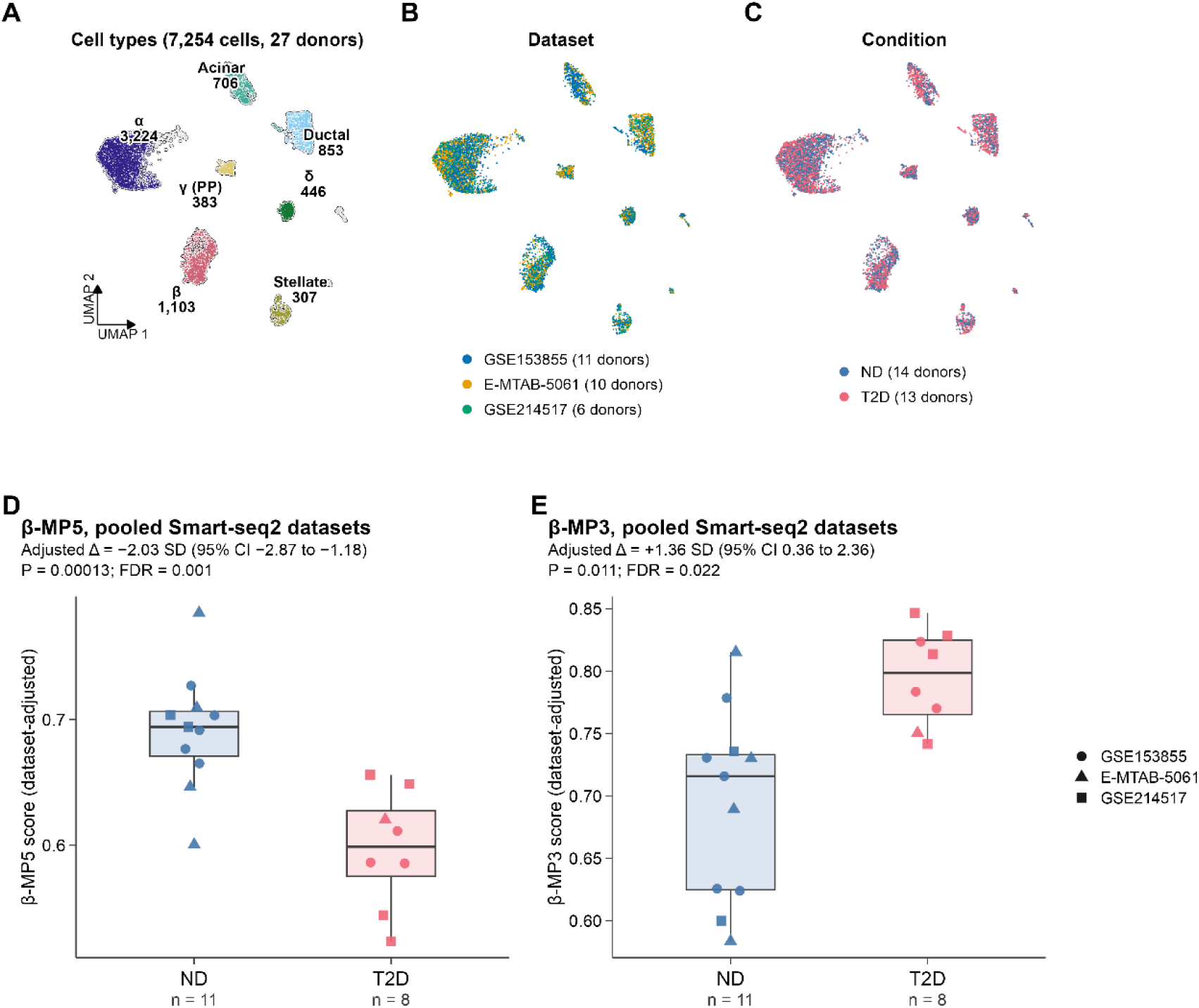
Secondary single-cell validation in pooled Smart-seq2 datasets. Three datasets (GSE153855, E-MTAB-5061 and GSE214517) were pooled for validation. (A–C) UMAP of 7,254 QC-passed cells from 27 donors, coloured by cell type (A), dataset (B) and diagnosis (C). (D, E) Donor-level β-MP5 and β-MP3 scores in the primary analysis population (11 ND and eight T2D donors). Scores were calculated using the frozen gene signatures and the shared gene background across the three datasets.

**Supplementary Figure 10.**
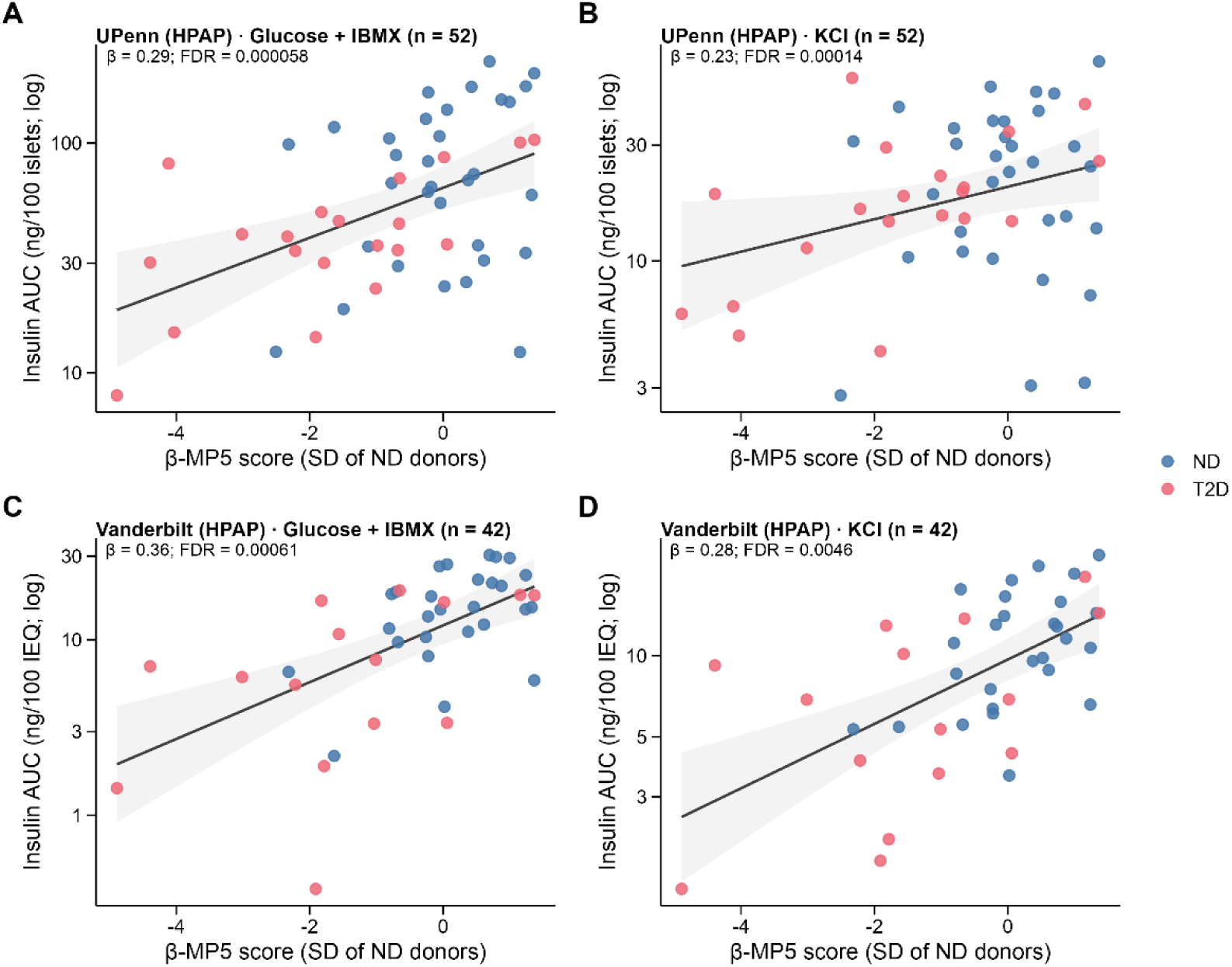
Additional associations between β-MP5 and stimulated insulin secretion. (A–D) Associations between β-MP5 scores and insulin AUC during glucose plus IBMX and KCl stimulation in UPenn (A, B) and Vanderbilt (C, D). Each point represents one donor. Insulin AUC is shown on a logarithmic scale. Lines indicate unadjusted fits; β estimates and FDR values are from the adjusted Model 1 regression.

**Supplementary Figure 11.**
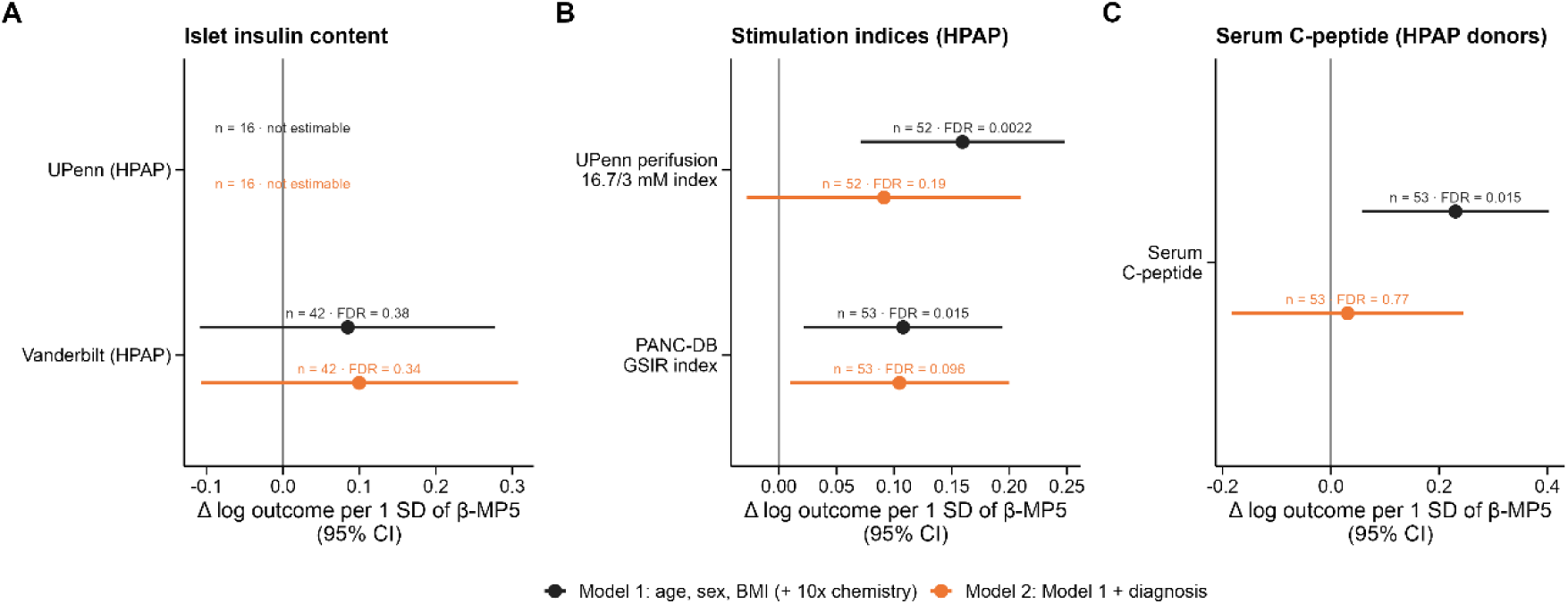
Associations between β-MP5 and additional functional measures. Adjusted effects per 1-SD increase in β-MP5 score under Model 1 (age, sex, BMI and 10x chemistry) and Model 2 (additionally adjusted for diagnosis). (A) Islet insulin content; the UPenn association was not estimable because only 16 donors had measurements. (B) UPenn glucose stimulation index and PANC-DB glucose-stimulated insulin response (GSIR) index. (C) Serum C-peptide in HPAP donors. Points indicate effect estimates and horizontal lines indicate 95% confidence intervals.

**Supplementary Figure 12.**
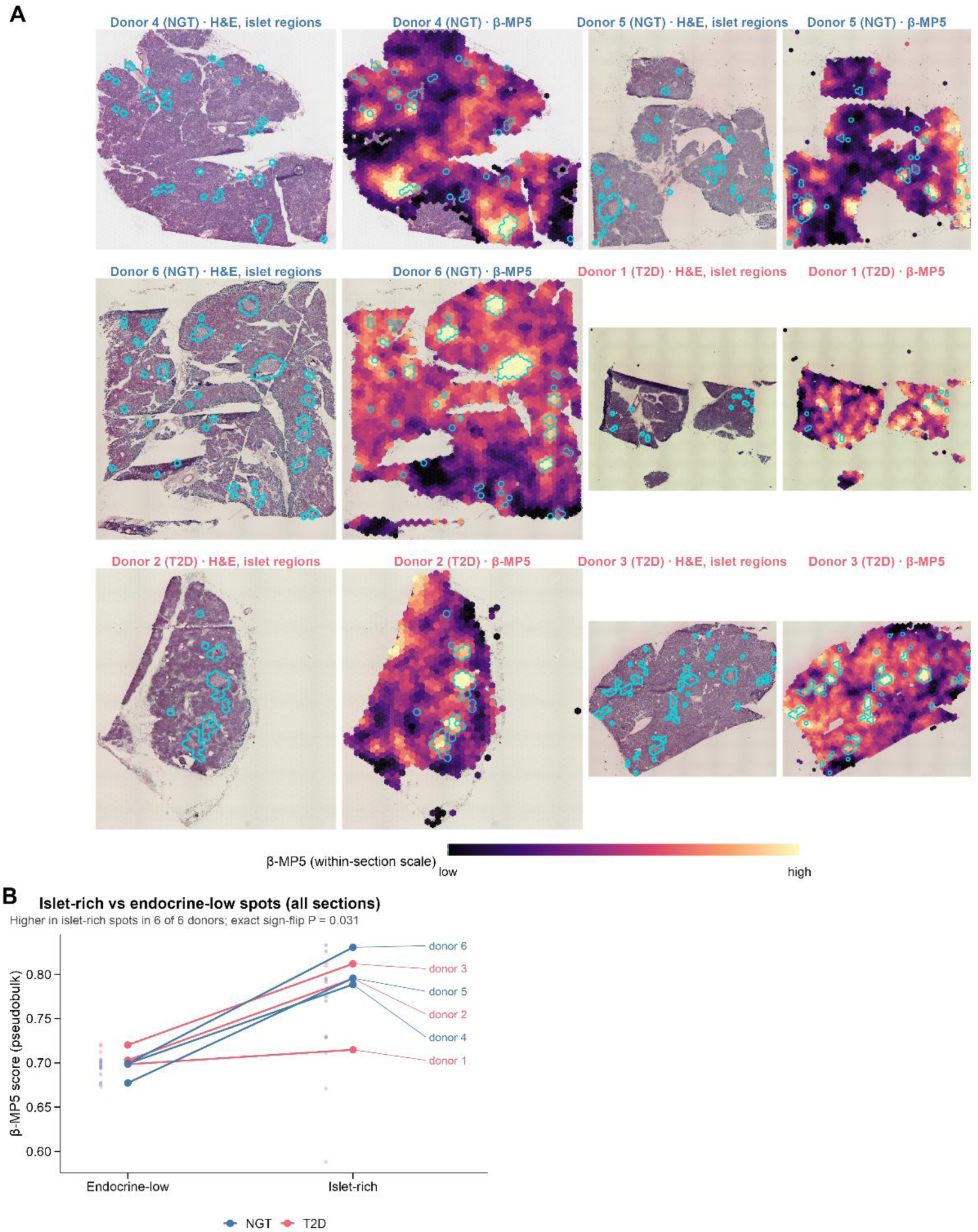
Spatial distribution of β-MP5 in human pancreas sections (GSE264331). Spatial transcriptomic profiles from 16 Visium sections from six donors (three NGT and three T2D) were analysed. Spots with fewer than 500 UMIs were excluded. Islet-rich spots were identified using seven non-hormone endocrine markers, none of which belonged to β-MP5. Intermediate spots were excluded, and spots at or below the expected marker count were classified as endocrine-low. (A) Representative section from each donor. Left, H&E image with transcriptomically defined islet regions outlined in cyan; right, spatial distribution of β-MP5 scores. Scores were smoothed across adjacent spots, and colour scales were clipped within each section; colours are not directly comparable between sections. (B) Donor-level pseudobulk β-MP5 scores in islet-rich and endocrine-low compartments using all available sections. Lines connect the two compartments within each donor. β-MP5 scores were higher in islet-rich spots in all six donors (exact two-sided sign-flip test, P = 0.031). Donors, rather than spots or sections, were the units of inference. NGT and T2D groups were not compared because of the small number of donors per group.

## Notes

### Competing Interest Statement

The authors have declared no competing interest.

## Reference

1. Sun, H., Saeedi, P., Karuranga, S., Pinkepank, M., Ogurtsova, K., Duncan, B.B., Stein, C., Basit, A., Chan, J.C.N., Mbanya, J.C., et al. (2022). IDF Diabetes Atlas: Global, regional and country-level diabetes prevalence estimates for 2021 and projections for 2045. Diabetes Res Clin Pract 183, 109119. 10.1016/j.diabres.2021.109119.

2. Kahn, S.E., Cooper, M.E., and Del Prato, S. (2014). Pathophysiology and treatment of type 2 diabetes: perspectives on the past, present, and future. Lancet 383, 1068–1083. 10.1016/S0140-6736(13)62154-6.

3. Prentki, M., and Nolan, C.J. (2006). Islet beta cell failure in type 2 diabetes. J Clin Invest 116, 1802–1812. 10.1172/JCI29103.

4. Butler, A.E., Janson, J., Bonner-Weir, S., Ritzel, R., Rizza, R.A., and Butler, P.C. (2003). Beta-cell deficit and increased beta-cell apoptosis in humans with type 2 diabetes. Diabetes 52, 102–110. 10.2337/diabetes.52.1.102.

5. Rahier, J., Guiot, Y., Goebbels, R.M., Sempoux, C., and Henquin, J.C. (2008). Pancreatic beta-cell mass in European subjects with type 2 diabetes. Diabetes Obes Metab 10 *Suppl 4*, 32–42. 10.1111/j.1463-1326.2008.00969.x.

6. Weir, G.C., Gaglia, J., and Bonner-Weir, S. (2020). Inadequate beta-cell mass is essential for the pathogenesis of type 2 diabetes. Lancet Diabetes Endocrinol 8, 249–256. 10.1016/S2213-8587(20)30022-X.

7. Del Guerra, S., Lupi, R., Marselli, L., Masini, M., Bugliani, M., Sbrana, S., Torri, S., Pollera, M., Boggi, U., Mosca, F., et al. (2005). Functional and molecular defects of pancreatic islets in human type 2 diabetes. Diabetes 54, 727–735. 10.2337/diabetes.54.3.727.

8. Talchai, C., Xuan, S., Lin, H.V., Sussel, L., and Accili, D. (2012). Pancreatic beta cell dedifferentiation as a mechanism of diabetic beta cell failure. Cell 150, 1223–1234. 10.1016/j.cell.2012.07.029.

9. Guo, S., Dai, C., Guo, M., Taylor, B., Harmon, J.S., Sander, M., Robertson, R.P., Powers, A.C., and Stein, R. (2013). Inactivation of specific beta cell transcription factors in type 2 diabetes. J Clin Invest 123, 3305–3316. 10.1172/JCI65390.

10. Cinti, F., Bouchi, R., Kim-Muller, J.Y., Ohmura, Y., Sandoval, P.R., Masini, M., Marselli, L., Suleiman, M., Ratner, L.E., Marchetti, P., and Accili, D. (2016). Evidence of beta-Cell Dedifferentiation in Human Type 2 Diabetes. J Clin Endocrinol Metab 101, 1044–1054. 10.1210/jc.2015-2860.

11. Lytrivi, M., Castell, A.L., Poitout, V., and Cnop, M. (2020). Recent Insights Into Mechanisms of beta-Cell Lipo- and Glucolipotoxicity in Type 2 Diabetes. J Mol Biol 432, 1514–1534. 10.1016/j.jmb.2019.09.016.

12. Gerber, P.A., and Rutter, G.A. (2017). The Role of Oxidative Stress and Hypoxia in Pancreatic Beta-Cell Dysfunction in Diabetes Mellitus. Antioxid Redox Signal 26, 501–518. 10.1089/ars.2016.6755.

13. Hudish, L.I., Reusch, J.E., and Sussel, L. (2019). beta Cell dysfunction during progression of metabolic syndrome to type 2 diabetes. J Clin Invest 129, 4001–4008. 10.1172/JCI129188.

14. Taylor, R., Al-Mrabeh, A., and Sattar, N. (2019). Understanding the mechanisms of reversal of type 2 diabetes. Lancet Diabetes Endocrinol 7, 726–736. 10.1016/S2213-8587(19)30076-2.

15. Dominguez, V., Raimondi, C., Somanath, S., Bugliani, M., Loder, M.K., Edling, C.E., Divecha, N., da Silva-Xavier, G., Marselli, L., Persaud, S.J., et al. (2011). Class II phosphoinositide 3-kinase regulates exocytosis of insulin granules in pancreatic beta cells. J Biol Chem 286, 4216–4225. 10.1074/jbc.M110.200295.

16. Fadista, J., Vikman, P., Laakso, E.O., Mollet, I.G., Esguerra, J.L., Taneera, J., Storm, P., Osmark, P., Ladenvall, C., Prasad, R.B., et al. (2014). Global genomic and transcriptomic analysis of human pancreatic islets reveals novel genes influencing glucose metabolism. Proc Natl Acad Sci U S A 111, 13924–13929. 10.1073/pnas.1402665111.

17. Solimena, M., Schulte, A.M., Marselli, L., Ehehalt, F., Richter, D., Kleeberg, M., Mziaut, H., Knoch, K.P., Parnis, J., Bugliani, M., et al. (2018). Systems biology of the IMIDIA biobank from organ donors and pancreatectomised patients defines a novel transcriptomic signature of islets from individuals with type 2 diabetes. Diabetologia 61, 641–657. 10.1007/s00125-017-4500-3.

18. Wigger, L., Barovic, M., Brunner, A.D., Marzetta, F., Schoniger, E., Mehl, F., Kipke, N., Friedland, D., Burdet, F., Kessler, C., et al. (2021). Multi-omics profiling of living human pancreatic islet donors reveals heterogeneous beta cell trajectories towards type 2 diabetes. Nat Metab 3, 1017–1031. 10.1038/s42255-021-00420-9.

19. Kayton, N.S., Poffenberger, G., Henske, J., Dai, C., Thompson, C., Aramandla, R., Shostak, A., Nicholson, W., Brissova, M., Bush, W.S., and Powers, A.C. (2015). Human islet preparations distributed for research exhibit a variety of insulin-secretory profiles. Am J Physiol Endocrinol Metab 308, E592–602. 10.1152/ajpendo.00437.2014.

20. Gloyn, A.L., Ibberson, M., Marchetti, P., Powers, A.C., Rorsman, P., Sander, M., and Solimena, M. (2022). Every islet matters: improving the impact of human islet research. Nat Metab 4, 970–977. 10.1038/s42255-022-00607-8.

21. Baron, M., Veres, A., Wolock, S.L., Faust, A.L., Gaujoux, R., Vetere, A., Ryu, J.H., Wagner, B.K., Shen-Orr, S.S., Klein, A.M., et al. (2016). A Single-Cell Transcriptomic Map of the Human and Mouse Pancreas Reveals Inter- and Intra-cell Population Structure. Cell Syst 3, 346–360 e344. 10.1016/j.cels.2016.08.011.

22. Muraro, M.J., Dharmadhikari, G., Grun, D., Groen, N., Dielen, T., Jansen, E., van Gurp, L., Engelse, M.A., Carlotti, F., de Koning, E.J., and van Oudenaarden, A. (2016). A Single-Cell Transcriptome Atlas of the Human Pancreas. Cell Syst 3, 385–394 e383. 10.1016/j.cels.2016.09.002.

23. Segerstolpe, A., Palasantza, A., Eliasson, P., Andersson, E.M., Andreasson, A.C., Sun, X., Picelli, S., Sabirsh, A., Clausen, M., Bjursell, M.K., et al. (2016). Single-Cell Transcriptome Profiling of Human Pancreatic Islets in Health and Type 2 Diabetes. Cell Metab 24, 593–607. 10.1016/j.cmet.2016.08.020.

24. Xin, Y., Kim, J., Okamoto, H., Ni, M., Wei, Y., Adler, C., Murphy, A.J., Yancopoulos, G.D., Lin, C., and Gromada, J. (2016). RNA Sequencing of Single Human Islet Cells Reveals Type 2 Diabetes Genes. Cell Metab 24, 608–615. 10.1016/j.cmet.2016.08.018.

25. Bader, E., Migliorini, A., Gegg, M., Moruzzi, N., Gerdes, J., Roscioni, S.S., Bakhti, M., Brandl, E., Irmler, M., Beckers, J., et al. (2016). Identification of proliferative and mature beta-cells in the islets of Langerhans. Nature 535, 430–434. 10.1038/nature18624.

26. Lawlor, N., George, J., Bolisetty, M., Kursawe, R., Sun, L., Sivakamasundari, V., Kycia, I., Robson, P., and Stitzel, M.L. (2017). Single-cell transcriptomes identify human islet cell signatures and reveal cell-type-specific expression changes in type 2 diabetes. Genome Res 27, 208–222. 10.1101/gr.212720.116.

27. Elgamal, R.M., Kudtarkar, P., Melton, R.L., Mummey, H.M., Benaglio, P., Okino, M.L., and Gaulton, K.J. (2023). An Integrated Map of Cell Type-Specific Gene Expression in Pancreatic Islets. Diabetes 72, 1719–1728. 10.2337/db23-0130.

28. Wang, X., Park, J., Susztak, K., Zhang, N.R., and Li, M. (2019). Bulk tissue cell type deconvolution with multi-subject single-cell expression reference. Nat Commun 10, 380. 10.1038/s41467-018-08023-x.

29. Mawla, A.M., and Huising, M.O. (2019). Navigating the Depths and Avoiding the Shallows of Pancreatic Islet Cell Transcriptomes. Diabetes 68, 1380–1393. 10.2337/dbi18-0019.

30. Bosi, E., Marselli, L., De Luca, C., Suleiman, M., Tesi, M., Ibberson, M., Eizirik, D.L., Cnop, M., and Marchetti, P. (2020). Integration of single-cell datasets reveals novel transcriptomic signatures of beta-cells in human type 2 diabetes. NAR Genom Bioinform 2, lqaa097. 10.1093/nargab/lqaa097.

31. Ngara, M., and Wierup, N. (2022). Lessons from single-cell RNA sequencing of human islets. Diabetologia 65, 1241–1250. 10.1007/s00125-022-05699-1.

32. Squair, J.W., Gautier, M., Kathe, C., Anderson, M.A., James, N.D., Hutson, T.H., Hudelle, R., Qaiser, T., Matson, K.J.E., Barraud, Q., et al. (2021). Confronting false discoveries in single-cell differential expression. Nat Commun 12, 5692. 10.1038/s41467-021-25960-2.

33. Crowell, H.L., Soneson, C., Germain, P.L., Calini, D., Collin, L., Raposo, C., Malhotra, D., and Robinson, M.D. (2020). muscat detects subpopulation-specific state transitions from multi-sample multi-condition single-cell transcriptomics data. Nat Commun 11, 6077. 10.1038/s41467-020-19894-4.

34. Kaestner, K.H., Powers, A.C., Naji, A., Consortium, H., and Atkinson, M.A. (2019). NIH Initiative to Improve Understanding of the Pancreas, Islet, and Autoimmunity in Type 1 Diabetes: The Human Pancreas Analysis Program (HPAP). Diabetes 68, 1394–1402. 10.2337/db19-0058.

35. Shapira, S.N., Naji, A., Atkinson, M.A., Powers, A.C., and Kaestner, K.H. (2022). Understanding islet dysfunction in type 2 diabetes through multidimensional pancreatic phenotyping: The Human Pancreas Analysis Program. Cell Metab 34, 1906–1913. 10.1016/j.cmet.2022.09.013.

36. Wang, G., Chiou, J., Zeng, C., Miller, M., Matta, I., Han, J.Y., Kadakia, N., Okino, M.L., Beebe, E., Mallick, M., et al. (2023). Integrating genetics with single-cell multiomic measurements across disease states identifies mechanisms of beta cell dysfunction in type 2 diabetes. Nat Genet 55, 984–994. 10.1038/s41588-023-01397-9.

37. Bandesh, K., Motakis, E., Nargund, S., Kursawe, R., Selvam, V., Ansarullah, Bhuiyan, R.M., Eryilmaz, G.N., Willett, A.M., White, J.K., et al. (2026). Deep single-cell decoding of human pancreatic islets reveals T2D beta-cell gene expression defects. EMBO J 45, 3978–4005. 10.1038/s44318-026-00744-w.

38. Martinez-Lopez, J.A., Lindqvist, A., Lopez-Pascual, A., Harder, A., Chen, P., Ngara, M., Shcherbina, L., Siffo, S., Cowan, E., Baira, S.M., et al. (2025). Single-cell mRNA-regulation analysis reveals cell type-specific mechanisms of type 2 diabetes. Nat Commun 16, 9475. 10.1038/s41467-025-65060-z.

39. Dorrell, C., Schug, J., Canaday, P.S., Russ, H.A., Tarlow, B.D., Grompe, M.T., Horton, T., Hebrok, M., Streeter, P.R., Kaestner, K.H., and Grompe, M. (2016). Human islets contain four distinct subtypes of beta cells. Nat Commun 7, 11756. 10.1038/ncomms11756.

40. Benninger, R.K.P., and Hodson, D.J. (2018). New Understanding of beta-Cell Heterogeneity and In Situ Islet Function. Diabetes 67, 537–547. 10.2337/dbi17-0040.

41. Avrahami, D., Wang, Y.J., Schug, J., Feleke, E., Gao, L., Liu, C., Consortium, H., Naji, A., Glaser, B., and Kaestner, K.H. (2020). Single-cell transcriptomics of human islet ontogeny defines the molecular basis of beta-cell dedifferentiation in T2D. Mol Metab 42, 101057. 10.1016/j.molmet.2020.101057.

42. Lee, D.D., and Seung, H.S. (1999). Learning the parts of objects by non-negative matrix factorization. Nature 401, 788–791. 10.1038/44565.

43. Kotliar, D., Veres, A., Nagy, M.A., Tabrizi, S., Hodis, E., Melton, D.A., and Sabeti, P.C. (2019). Identifying gene expression programs of cell-type identity and cellular activity with single-cell RNA-Seq. Elife 8, e43803. 10.7554/eLife.43803.

44. Kinker, G.S., Greenwald, A.C., Tal, R., Orlova, Z., Cuoco, M.S., McFarland, J.M., Warren, A., Rodman, C., Roth, J.A., Bender, S.A., et al. (2020). Pan-cancer single-cell RNA-seq identifies recurring programs of cellular heterogeneity. Nat Genet 52, 1208–1218. 10.1038/s41588-020-00726-6.

45. Barkley, D., Moncada, R., Pour, M., Liberman, D.A., Dryg, I., Werba, G., Wang, W., Baron, M., Rao, A., Xia, B., et al. (2022). Cancer cell states recur across tumor types and form specific interactions with the tumor microenvironment. Nat Genet 54, 1192–1201. 10.1038/s41588-022-01141-9.

46. Gavish, A., Tyler, M., Greenwald, A.C., Hoefflin, R., Simkin, D., Tschernichovsky, R., Galili Darnell, N., Somech, E., Barbolin, C., Antman, T., et al. (2023). Hallmarks of transcriptional intratumour heterogeneity across a thousand tumours. Nature 618, 598–606. 10.1038/s41586-023-06130-4.

47. Zhang, C., Zhang, Y., Wu, Z., Zhang, Y., Xue, F., Tan, Q., Zhong, X., Zhang, Y., Zhao, Z., Peng, Y., et al. (2026). Screening of cell-type-specific meta-programs for drug repurposing in Alzheimer’s disease. Brief Bioinform 27, bbag411. 10.1093/bib/bbag411.

48. Camunas-Soler, J., Dai, X.Q., Hang, Y., Bautista, A., Lyon, J., Suzuki, K., Kim, S.K., Quake, S.R., and MacDonald, P.E. (2020). Patch-Seq Links Single-Cell Transcriptomes to Human Islet Dysfunction in Diabetes. Cell Metab 31, 1017–1031 e1014. 10.1016/j.cmet.2020.04.005.

49. Howell, N., Weiss, Z., Bonnycastle, L.L., Grenko, C.M., Randazzo, D., Dampier, C.H., Sinha, N., Narisu, N., Swift, A.J., Erdos, M.R., et al. (2025). A spatial transcriptomics dataset of pancreas sections in normal glucose tolerance and type 2 diabetic donors. Sci Data 12, 1526. 10.1038/s41597-025-05450-6.

50. Germain, P.L., Lun, A., Garcia Meixide, C., Macnair, W., and Robinson, M.D. (2021). Doublet identification in single-cell sequencing data using scDblFinder. F1000Res 10, 979. 10.12688/f1000research.73600.2.

51. Hao, Y., Stuart, T., Kowalski, M.H., Choudhary, S., Hoffman, P., Hartman, A., Srivastava, A., Molla, G., Madad, S., Fernandez-Granda, C., and Satija, R. (2024). Dictionary learning for integrative, multimodal and scalable single-cell analysis. Nat Biotechnol 42, 293–304. 10.1038/s41587-023-01767-y.

52. Korsunsky, I., Millard, N., Fan, J., Slowikowski, K., Zhang, F., Wei, K., Baglaenko, Y., Brenner, M., Loh, P.R., and Raychaudhuri, S. (2019). Fast, sensitive and accurate integration of single-cell data with Harmony. Nat Methods 16, 1289–1296. 10.1038/s41592-019-0619-0.

53. Becht, E., McInnes, L., Healy, J., Dutertre, C.A., Kwok, I.W.H., Ng, L.G., Ginhoux, F., and Newell, E.W. (2018). Dimensionality reduction for visualizing single-cell data using UMAP. Nat Biotechnol 37, 38–44. 10.1038/nbt.4314.

54. Andreatta, M., and Carmona, S.J. (2021). UCell: Robust and scalable single-cell gene signature scoring. Comput Struct Biotechnol J 19, 3796–3798. 10.1016/j.csbj.2021.06.043.

55. Gaujoux, R., and Seoighe, C. (2010). A flexible R package for nonnegative matrix factorization. BMC Bioinformatics 11, 367. 10.1186/1471-2105-11-367.

56. Liberzon, A., Birger, C., Thorvaldsdottir, H., Ghandi, M., Mesirov, J.P., and Tamayo, P. (2015). The Molecular Signatures Database (MSigDB) hallmark gene set collection. Cell Syst 1, 417–425. 10.1016/j.cels.2015.12.004.

57. Kanehisa, M., and Goto, S. (2000). KEGG: kyoto encyclopedia of genes and genomes. Nucleic Acids Res 28, 27–30. 10.1093/nar/28.1.27.

58. Ashburner, M., Ball, C.A., Blake, J.A., Botstein, D., Butler, H., Cherry, J.M., Davis, A.P., Dolinski, K., Dwight, S.S., Eppig, J.T., et al. (2000). Gene ontology: tool for the unification of biology. The Gene Ontology Consortium. Nat Genet 25, 25–29. 10.1038/75556.

59. Wu, T., Hu, E., Xu, S., Chen, M., Guo, P., Dai, Z., Feng, T., Zhou, L., Tang, W., Zhan, L., et al. (2021). clusterProfiler 4.0: A universal enrichment tool for interpreting omics data. Innovation (Camb) 2, 100141. 10.1016/j.xinn.2021.100141.

60. Benjamini, Y., and Hochberg, Y. (1995). Controlling the False Discovery Rate - a Practical and Powerful Approach to Multiple Testing. Journal of the Royal Statistical Society Series B-Statistical Methodology 57, 289–300. DOI 10.1111/j.2517-6161.1995.tb02031.x.

61. Robinson, M.D., and Oshlack, A. (2010). A scaling normalization method for differential expression analysis of RNA-seq data. Genome Biol 11, R25. 10.1186/gb-2010-11-3-r25.

62. Mackinnon, J.G., and White, H. (1985). Some Heteroskedasticity-Consistent Covariance-Matrix Estimators with Improved Finite-Sample Properties. Journal of Econometrics 29, 305–325. Doi 10.1016/0304-4076(85)90158-7.

63. Long, J.S., and Ervin, L.H. (2000). Using heteroscedasticity consistent standard errors in the linear regression model. American Statistician 54, 217–224. Doi 10.2307/2685594.

64. Zeileis, A. (2004). Econometric Computing with HC and HAC Covariance Matrix Estimators. Journal of Statistical Software 11. 10.18637/jss.v011.i10.

65. Chen, Y., Lun, A.T., and Smyth, G.K. (2016). From reads to genes to pathways: differential expression analysis of RNA-Seq experiments using Rsubread and the edgeR quasi-likelihood pipeline. F1000Res 5, 1438. 10.12688/f1000research.8987.2.

66. Irizarry, R.A., Hobbs, B., Collin, F., Beazer-Barclay, Y.D., Antonellis, K.J., Scherf, U., and Speed, T.P. (2003). Exploration, normalization, and summaries of high density oligonucleotide array probe level data. Biostatistics 4, 249–264. 10.1093/biostatistics/4.2.249.

67. Ritchie, M.E., Phipson, B., Wu, D., Hu, Y., Law, C.W., Shi, W., and Smyth, G.K. (2015). limma powers differential expression analyses for RNA-sequencing and microarray studies. Nucleic Acids Res 43, e47. 10.1093/nar/gkv007.

68. Chu, T., Wang, Z., Pe’er, D., and Danko, C.G. (2022). Cell type and gene expression deconvolution with BayesPrism enables Bayesian integrative analysis across bulk and single-cell RNA sequencing in oncology. Nat Cancer 3, 505–517. 10.1038/s43018-022-00356-3.

69. Newman, A.M., Steen, C.B., Liu, C.L., Gentles, A.J., Chaudhuri, A.A., Scherer, F., Khodadoust, M.S., Esfahani, M.S., Luca, B.A., Steiner, D., et al. (2019). Determining cell type abundance and expression from bulk tissues with digital cytometry. Nat Biotechnol 37, 773–782. 10.1038/s41587-019-0114-2.

70. Cataldo, L.R., Vishnu, N., Singh, T., Bertonnier-Brouty, L., Bsharat, S., Luan, C., Renstrom, E., Prasad, R.B., Fex, M., Mulder, H., and Artner, I. (2021). The MafA-target gene PPP1R1A regulates GLP1R-mediated amplification of glucose-stimulated insulin secretion in beta-cells. Metabolism 118, 154734. 10.1016/j.metabol.2021.154734.

71. Carrat, G.R., Hu, M., Nguyen-Tu, M.S., Chabosseau, P., Gaulton, K.J., van de Bunt, M., Siddiq, A., Falchi, M., Thurner, M., Canouil, M., et al. (2017). Decreased STARD10 Expression Is Associated with Defective Insulin Secretion in Humans and Mice. Am J Hum Genet 100, 238–256. 10.1016/j.ajhg.2017.01.011.

72. Vierra, N.C., Dadi, P.K., Jeong, I., Dickerson, M., Powell, D.R., and Jacobson, D.A. (2015). Type 2 Diabetes-Associated K+ Channel TALK-1 Modulates beta-Cell Electrical Excitability, Second-Phase Insulin Secretion, and Glucose Homeostasis. Diabetes 64, 3818–3828. 10.2337/db15-0280.

73. Refai, E., Dekki, N., Yang, S.N., Imreh, G., Cabrera, O., Yu, L., Yang, G., Norgren, S., Rossner, S.M., Inverardi, L., et al. (2005). Transthyretin constitutes a functional component in pancreatic beta-cell stimulus-secretion coupling. Proc Natl Acad Sci U S A 102, 17020–17025. 10.1073/pnas.0503219102.

74. Flamez, D., Roland, I., Berton, A., Kutlu, B., Dufrane, D., Beckers, M.C., De Waele, E., Rooman, I., Bouwens, L., Clark, A., et al. (2010). A genomic-based approach identifies FXYD domain containing ion transport regulator 2 (FXYD2)gammaa as a pancreatic beta cell-specific biomarker. Diabetologia 53, 1372–1383. 10.1007/s00125-010-1714-z.

75. Avrahami, D., Li, C., Yu, M., Jiao, Y., Zhang, J., Naji, A., Ziaie, S., Glaser, B., and Kaestner, K.H. (2014). Targeting the cell cycle inhibitor p57Kip2 promotes adult human beta cell replication. J Clin Invest 124, 670–674. 10.1172/JCI69519.

76. Rorsman, P., and Ashcroft, F.M. (2018). Pancreatic beta-Cell Electrical Activity and Insulin Secretion: Of Mice and Men. Physiol Rev 98, 117–214. 10.1152/physrev.00008.2017.

77. Stahl, P.L., Salmen, F., Vickovic, S., Lundmark, A., Navarro, J.F., Magnusson, J., Giacomello, S., Asp, M., Westholm, J.O., Huss, M., et al. (2016). Visualization and analysis of gene expression in tissue sections by spatial transcriptomics. Science 353, 78–82. 10.1126/science.aaf2403.

78. Ahlqvist, E., Storm, P., Karajamaki, A., Martinell, M., Dorkhan, M., Carlsson, A., Vikman, P., Prasad, R.B., Aly, D.M., Almgren, P., et al. (2018). Novel subgroups of adult-onset diabetes and their association with outcomes: a data-driven cluster analysis of six variables. Lancet Diabetes Endocrinol 6, 361–369. 10.1016/S2213-8587(18)30051-2.

